# Understanding the Global Spread of Artemisinin Resistance: Insights from over 100K *Plasmodium falciparum* Samples

**DOI:** 10.1101/2024.08.23.609367

**Authors:** Andrew J. Balmer, Nina F. D. White, Eyyüb S. Ünlü, Chiyun Lee, Richard D. Pearson, Jacob Almagro-Garcia, Cristina V. Ariani

**Author notes:** These authors contributed equally to this work (Andrew J. Balmer, Nina F. D. White, Eyyüb S. Ünlü).

## Abstract

Artemisinin partial resistance (ART-R) in *Plasmodium falciparum* is one of the most pressing threats to global malaria control. Over the last two decades, ART-R has spread widely across Southeast Asia, compromising public health strategies and hindering elimination efforts. As of 2024, ART-R has now emerged in East Africa, with the potential to dramatically increase human mortality in the region. Mitigating the spread of ART-R requires detailed genomic surveillance of point mutations in the *kelch13* gene, the primary determinant of resistance to artemisinin derivatives. Although extensive surveillance data on these markers is available, it is distributed across many literature studies and open databases. In this literature review, we aggregate publicly available spatiotemporal data for 112,933 *P. falciparum* samples between 1980 – 2023 into a single resource, providing the most comprehensive overview of *kelch13* markers to date. By synthesising insights from these samples over a global scale, we outline the history and current status of *kelch13* mutations associated with ART-R, with particular reference to their emergence in Southeast Asia and recent emergence in East and Northeast Africa. Concerningly, we find their recent increases in frequency in these areas of Africa are comparable to those observed in Southeast Asia 10-15 years ago. We review several factors that may influence the spread of ART-R going forwards, such as fitness costs, treatment strategies, and local epidemiological dynamics, before articulating possible scenarios on how resistance may spread in Africa in coming years. In summary, this review provides a unified, comprehensive account of how the situation of ART-R has unfolded globally so far, highlighting insights both for researchers in the field and public health bodies which aim to reduce its negative effects. More broadly, we highlight the critical role genomic surveillance has had, and will continue to have in combating the spread of ART-R.

## 1. Introduction

### 1.1 Background to artemisinin partial resistance

By the 1990s, resistance to two major antimalarial drugs, chloroquine and sulfadoxine-pyrimethamine, was widespread across malaria-endemic countries (Wellems and Plowe, 2001; Gatton, Martin and Cheng, 2004). In the following decade, these drugs were replaced by the now widely used artemisinin-combination therapies (ACTs), which consist of a fast-acting artemisinin derivative and a long-acting partner drug. ACTs are well-tolerated and highly effective for treating *falciparum* malaria (Nosten and White, 2007), the most severe form of the disease. For these reasons, in 2006 the World Health Organisation (WHO) officially recommended ACTs as first-line treatments for uncomplicated *falciparum* malaria and intravenous artesunate for severe malaria (WHO, 2006). By 2022, ACTs had been adopted by 84 countries and was the most widely-used treatment (WHO, 2023). Despite the development of new drugs, the lengthy testing and clinical trial process means ACTs will remain the primary treatment for *P. falciparum* malaria for the foreseeable future (Erhunse and Sahal, 2021).

Reports of reduced efficacy of artemisinin-based drugs emerged from the Thai-Cambodian border in 2008. Noedl et al. (2008) reported prolonged parasite-clearance times (133 and 95 hours) in two patients, compared with a median of 52.2 hours for patients who were cured. Similarly, Dondorp et al. (2009) found that median parasite-clearance was prolonged by 6 hours in the Thai-Cambodian border group compared to the group from northwestern Thailand. These were the first recorded instances of what we now refer to as artemisinin partial resistance (hereafter referred to as ART-R). ART-R is characterised by a delayed parasite clearance time and is described as ‘partial’ resistance as only ring-stage parasites survive artemisinin (Behrens, Schmidt and Spielmann, 2021). Specifically, ART-R is determined following treatment of *falciparum* malaria with an artemisinin monotherapy or ACT if persistent parasitemia occurs after three days or if the half-life of parasite clearance from the bloodstream exceeds 5.5 hours (Rosenthal, 2021). Following the emergence of ART-R, it was identified across the Greater Mekong Subregion, in Myanmar, Vietnam, and Laos (Ashley *et al*., 2014; Tun *et al*., 2015).

Just as with the appearance of resistance to previous antimalarial drugs chloroquine and sulfadoxine–pyrimethamine (SP), Southeast Asia became the initial origin point of ART-R. Following the spread of chloroquine and SP resistance across malaria endemic countries, a major concern now is the spread of ART-R in Africa. Africa carries 94% of the global malaria burden (WHO, 2023) and ACTs are the predominant first-line defence against malaria there. Artemisinin and its derivatives have a short half-life within patients of two to five hours and in ACTs are responsible for the initial, rapid reduction of parasitemia (de Vries and Dien, 1996). ART-R poses a risk to ACT efficacy because delayed parasite clearance places a larger parasite burden on partner drugs and increases the chances of resistance to partner drugs developing. While ACTs remain largely effective in Africa, reports of reduced ACT efficacy have begun to emerge for different ACTs in Africa - artemether–lumefantrine (AL), dihydroartemisinin–piperaquine (DHA-PPQ) and artesunate–amodiaquine (ASAQ). Efficacy below 90% has been reported in the countries of Angola (AL) (Dimbu *et al*., 2021), the DRC (AL, DHA-PPQ) (Moriarty *et al*., 2021), Burkina Faso (AL, DHA-PPQ) (Gansané *et al*., 2021), Uganda (AL) (Ebong *et al*., 2021), and Tanzania (AL, ASAQ) (Ishengoma *et al*., 2024). If the efficacy of artemisinin derivatives should decline in tandem with a rise in resistance to ACT partner drugs in Africa, the consequences would be dire (Duffey *et al*., 2021).

In addition to the clinical presentation of delayed clearance, molecular markers in the *kelch13* gene can aid in diagnosing ART-R (Ariey *et al*., 2014). Endemic ART-R can be confirmed when ≥5% of patients have an ART-R validated mutation marker in *kelch13* and meet the clinical diagnosis for ART-R (WHO, 2022). Analysis of *kelch13* in an *in vitro* engineered artemisinin-resistant parasite line and in field isolates collected from Cambodia revealed four mutations, each within a kelch repeat of the C-terminal *kelch13* propeller domain, linked with significantly higher ring-stage survival rates compared with the wild type (Ariey *et al*., 2014). These mutations were C580Y, R539T, Y493H, and I543T. Additional evidence for the association between *kelch13* mutations and ART-R was provided in a gene editing experiment (Straimer *et al*., 2015). Removal of *kelch13* mutations reduced parasite survival rates from 49% to 2.4%, and insertion of mutations into wild type isolates increased their survival from 0.6% to 29% (Straimer *et al*., 2015). Further studies, including genome-wide association analysis (Cerqueira *et al*., 2017) and scans for selection (Anderson *et al*., 2017) have contributed to our understanding of the role of *kelch13* in conferring ART-R.

Today, the WHO maintains a list of *kelch13* mutations linked to ART-R (WHO, 2022). The criteria for qualification as a candidate or validated marker of ART-R are shown in Table S1. Currently, the list contains 13 validated mutations and ten candidate mutations. All current mutations are in the BTB/POZ or propeller domain of *kelch13*, though a mutation outside this domain, E252Q, has been shown to be associated with delayed parasite clearance (Phyo *et al*., 2016; Amaratunga *et al*., 2019). This association appears to occur only in the presence of other mutations based on *in vitro* work, however (WHO, 2022). At least one other *kelch13* mutation in the BTB/POZ domain, P413A, has been associated with ART-R (Paloque *et al*., 2022), but has not yet met the criteria to be considered a validated/candidate marker by the WHO (Table S1).

### 1.2 Monitoring ART-R

ART-R is primarily monitored using three separate methods, each of which have their own costs and benefits (Nsanzabana *et al*., 2018). These methods include therapeutic efficacy studies (TES), *in vitro* assays of phenotypic resistance, and the genotyping of parasites to identify molecular markers associated with resistance (Nsanzabana *et al*., 2018). TES, recognised as the ‘gold-standard’ for monitoring treatment efficacy, are randomised clinical trials that assess treatment outcomes *in-vivo* over the course of several weeks and have been a major determinant in shaping antimalarial policies (WHO, 2009). For *in-vitro* assays of phenotypic resistance, parasites are directly exposed to varying concentrations of antimalarials to assess their susceptibility (WHO, 2009).

Alongside these methods, genomic epidemiology and surveillance has emerged as an additional strategy for monitoring ART-R. Genomic surveillance of molecular markers is a scalable method of monitoring the prevalence of resistance at population levels (Ndiaye *et al*., 2021). As well as identifying resistance-associated variants, genomic surveillance can track accompanying signs of parasite adaptations to control measures, such as clonal population expansions, runs of extended homozygosity, and increases in copy number variation (Duffy *et al*., 2017; Jacob *et al*., 2021). Genomic surveillance has been increasingly implemented in malaria endemic countries and has the potential to reduce the time between sample collection and data availability, providing earlier warnings for planning public policy strategies. Although genomic surveillance methods do not directly measure effects on patient outcome, they can be particularly informative when combined with results of therapeutic efficacy or *in vitro* work (Yeka *et al*., 2019; Ippolito *et al*., 2020). For example, GWAS studies have revealed the role of various mutants in generating resistance, allowing resistance markers to be tracked at population scales (Miotto *et al*., 2015). Because genomic surveillance has wide scale applicability, there are now several strategies available to monitor molecular markers associated with ART-R over time and across geographical areas, including targeted amplicon and whole genome sequencing, as reviewed in (Neafsey, Taylor and MacInnis, 2021). These strategies have made it possible to collect wide-scale and detailed datasets on prevalence of important molecular markers at population scales (MalariaGEN, 2023). For example, Flegg et al. (Flegg *et al*., 2024) have demonstrated the spatial and temporal trends in the spread of ART-R in Southeast Asia and Ndwiga et al. (2021) in Africa. Both studies showed that genomic surveillance of *kelch13* is a fast and effective method for measuring ART-R, though both focused on just one geographic region.

### 1.3 Review Rationale and Approach

Understanding how, where, and why ART-R occurs is essential for developing malaria control strategies. In this review, we provide a global perspective on the emergence, spread, and predicted next stages of ART-R. We collate publicly available data from over 112,000 samples, providing the most comprehensive overview of *kelch13* mutations to date. Our global-scale analysis of this reliable marker of ART-R allows us to provide a thorough overview of the history and current state of ART-R. We point to key spatio-temporal trends in the spread of ART-R in regions of Southeast Asia and East Africa. Building on these novel analyses and existing literature, we articulate possible scenarios for the next stage of ART-R in Africa. Finally, we discuss the role of genomic surveillance in effective malaria control moving forwards.

## 2. Spatiotemporal analysis of *kelch13* in 112,933 samples

### 2.1 Data Collection

We aggregated publicly available *kelch13* mutation data for 112,933 samples collected between 1980 – 2023, across 73 countries (Table S2, Supp. Dataset S1). This data and associated metadata was gathered from the MalariaGEN Pf7 release (MalariaGEN et al. 2023), the Worldwide Antimalarial Resistance Network (WWARN) Artemisinin Resistance Molecular Surveyor (www.iddo.org/wwarn/tracking-resistance/artemisinin-molecular-surveyor), and our own literature search (Supplementary Methods). We identified mutations which have been validated for an association with ART-R or marked as a candidate marker of ART-R according to the WHO’s guidelines (Table S1). We assigned each country to one of thirteen broader populations (Supplementary Methods, Table S2): South America, West Africa, Central Africa, North Africa, Northeast Africa, East Africa, Southern Africa, Western Asia, Eastern South Asia, Far-Eastern South Asia, Western Southeast Asia, Eastern Southeast Asia, and Oceania.

### 2.2 Global Geographic Overview

Across all populations, we observed 492 unique, non-synonymous mutations in the BTB/POZ and propeller domains of *kelch13,* alongside the 3D7 reference gene sequence (Supp. Dataset S2). We considered parasites with the 3D7 reference *kelch13* sequence and the mutation A578S to be susceptible to artemisinin (WHO, 2022). We hereafter refer to all BTB/POZ and propeller mutations (codons 349 - 726) as “propeller mutations”. The *kelch13* 3D7 reference sequence was present in the majority of samples (85.5%, n = 96,574), while the most common mutation was the validated marker C580Y at 6.5% (n = 7,307). The other most common mutations were all WHO validated/candidate: F446I (1.5%), R561H (0.7%), P441L (0.7%), and R539T (0.4%).

Of all populations, those in Southeast Asia had the highest proportions of samples with any *kelch13* propeller mutation and of samples with a WHO validated/candidate mutation (Figure 1A). In Eastern Southeast Asia, over half of samples (52%) had a *kelch13* propeller mutation and 98% of these mutant samples had a WHO validated/candidate marker. In Western Southeast Asia, 35% of samples were mutant at the *kelch13* propeller domain and 94% of these samples had a WHO validated/candidate mutation (Figure 1A). The ratio of WHO validated/candidate mutations relative to the total number of *kelch13* propeller mutations was also largest in Southeast Asia (Figure 1B). In Western Southeast Asia, this ratio was 0.19, while it was 0.32 in Eastern Southeast Asia. Furthermore, the biggest diversity of WHO validated/candidate markers out of all populations was found across these two Southeast Asian populations (Figure 1B).

**Figure 1.**
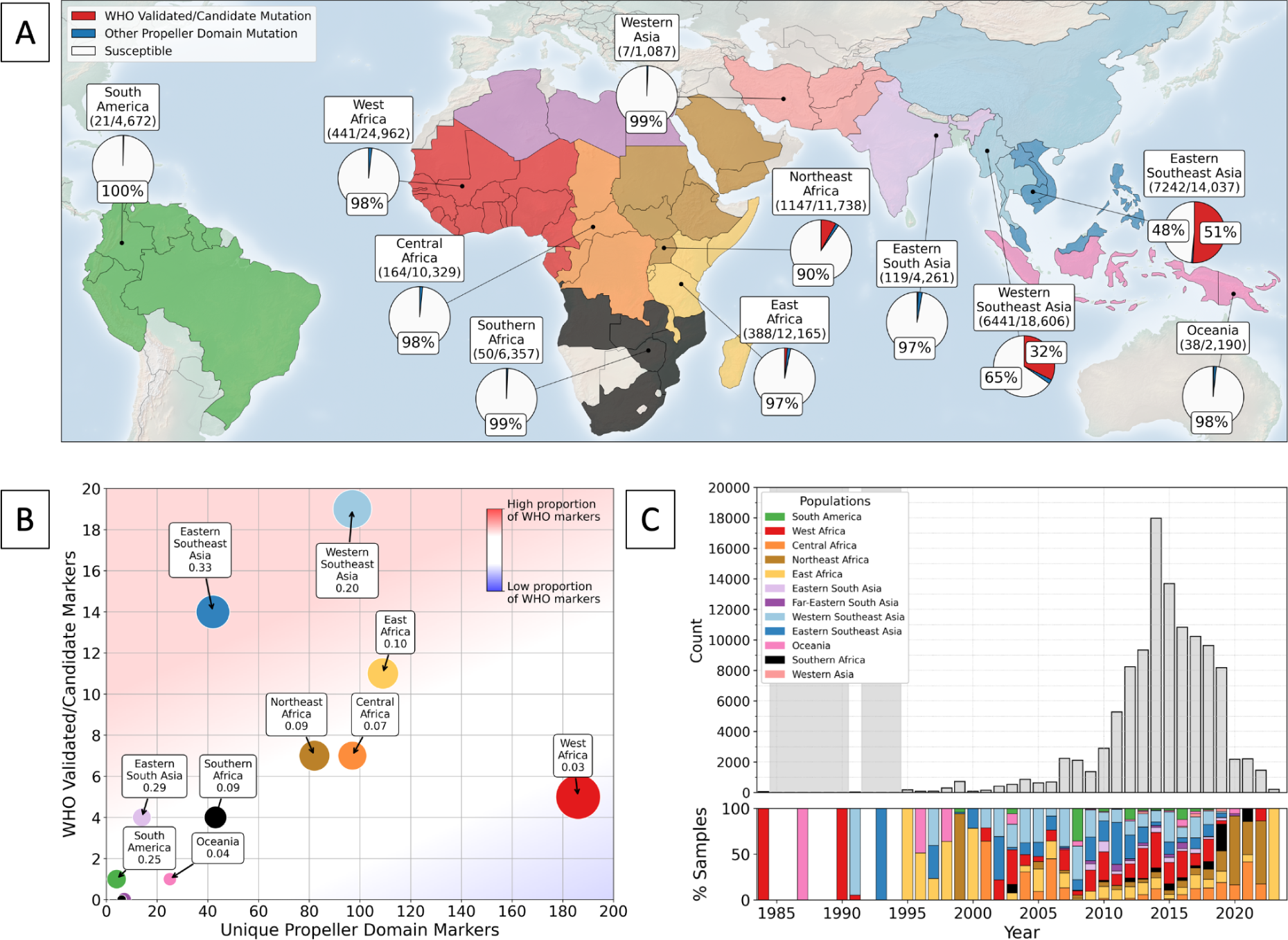
Global trends in *kelch13* propeller mutations over time. Panel A shows the distribution of *kelch13* mutations globally for samples collected between 1980-2023. Countries from which samples were included are coloured according to their geographic population assignment. All populations are labelled with a fraction, denoting the number of samples with any *kelch13* propeller mutation (excluding A578S, which is known to be artemisinin susceptible) over the total number of samples from each population. No mutations were detected in North African samples, therefore it is not labelled. For each population, pie charts show the proportion of samples with a WHO validated/candidate mutation in red, the proportion of any other propeller domain mutation in blue, and the proportion of samples with a 3D7 reference *kelch13* sequence or A578S mutation in white. Proportions of each category at 25% or more are labelled. Panel B shows the number of unique *kelch13* propeller domain markers for each population against their total number of unique WHO validated/candidate mutations. Populations are labelled with their ratio of WHO validated/candidate markers to the total number of unique propeller domain markers. Populations falling in the blue shaded area have fewer WHO validated/candidate markers relative to their total count of *kelch13* mutations than populations falling in red areas. The upper plot of panel C shows the total number of samples over time. Grey areas denote years with fewer than 25 samples. The bottom panel shows the proportion of samples from each population for each year, with years with no samples coloured white. Note the limited number of samples from Southeast Asia after 2019, which coincided with an increased proportion of resistant samples in East and Northeast Africa.

Africa was the next most prominent centre of high *kelch13* propeller mutation frequency and diversity after Southeast Asia (Figure 1A). In particular, populations in the east and northeast of the continent had higher proportions of samples with *kelch13* propeller mutations and with WHO validated/candidate markers, relative to the rest of Africa (Figure 1A). In Northeast Africa, 10% of all samples had a *kelch13* propeller mutation, ∼80% of which were WHO validated/candidate. In East Africa, 4% of samples had a *kelch13* propeller mutation, and a quarter of these were WHO validated/candidate markers for ART-R (Table S3). In other areas of Africa, the proportion of mutant samples was relatively low (Figure 1A). In West and Central Africa, mutant samples made up 2% of the total sample numbers and in Southern Africa this figure was 1%. The proportion of samples with a WHO validated/candidate mutation in West, Central, and Southern Africa was also lower than the eastern side of Africa, with proportions below 0.13% (Table S3). No *kelch13* propeller mutations were detected in North Africa, although only nine samples were collected here. These findings align with recent observations of rising ART-R in Eastern regions of Africa.

The east and northeast regions of Africa were also notable within Africa for their high ratios of WHO validated/candidate mutation diversity relative to their total numbers of *kelch13* propeller mutations (Figure 1B). In East Africa, the ratio of unique WHO validated/candidate mutations to total *kelch13* propeller mutations was 0.1 (i. e. 0.1 WHO mutation for every 1 unique propeller domain mutation). In Northeast Africa, this ratio was 0.09 (Figure 1B, Table S3). WHO validated/candidate mutation diversity was comparable between Northeast and Central Africa, with seven unique validated/candidate *kelch13* propeller mutations in each population (Figure 1B). The ratio to total *kelch13* propeller mutations was lower in Central Africa, however, at 0.07. Despite the fact that West Africa was the largest study population of all, with the largest number of unique *kelch13* propeller mutations (n = 185), only five mutations were WHO validated/candidate - a ratio of 0.03 (Table S3). This implies that there have not been strong pressures to evolve known WHO validated/candidate mutations in West Africa.

Outside of Southeast Asia and Africa, the rest of the world had lower mutant sample proportions and *kelch13* mutation diversity (Figure 1A). In South America, 0.45% of all samples held any *kelch13* propeller mutation and only three unique mutations have been reported in the region to date (Table S3). One WHO validated/candidate mutation was included in this count, found in 0.41% of all South American samples (Table S3). The second smallest population, Western Asia, contained only 0.64% mutant samples and five *kelch13* propeller mutations - none of which were WHO validated/candidate (Figure 1A). In Eastern South Asia, 3% of all samples had one of 13 *kelch13* propeller mutations detected in the population. Four WHO validated/candidate mutations were found in Eastern South Asia, which accounted for 0.54% of all samples there (Figure 1A). In Far-Eastern South Asia, levels of *kelch13* mutation diversity were lower. Only 0.52% of samples held any *kelch13* propeller mutations, and there were no WHO validated/candidate mutations recorded. In Oceania, ∼2% of all samples were mutant in the *kelch13* propeller domain (Figure 1A). Just one of the 24 total unique *kelch13* propeller mutations were WHO validated/candidate in Oceania (Figure 1A). For a complete list of mutation metrics in all populations, see Table S3.

### 2.3 Temporal Overview

Both sampling effort and the number of detected *kelch13* propeller mutations increased at a time when artemisinin monotherapies and ACTs became more commonly used (Figure 1C). Before 2000 there was limited sampling, with a mean of 138 samples genotyped per year globally (Table S4). Despite pre-2000s sampling occurring across four continents, samples predominantly had the *kelch13* 3D7 reference sequence during this time (Figure 1C). The first *kelch13* propeller polymorphisms were evident from 1991 and the first WHO validated/candidate mutation to be documented was R359T in Thailand, 1997. Another WHO validated/candidate mutation, V568G, was detected in Kenya at some point between 1996 and 2003, but exact years were not provided by the study (de Laurent *et al*., 2018). No other markers of ART-R were detected before 2000. After 2000, sample numbers increased, reaching up to 17,980 in 2014 (Figure 1C, Table S4). Pooling data from all populations, the global number of unique *kelch13* mutations and validated/candidate mutations both rose throughout the 2000s alongside the rise in annual sample size (Figure S1). Unique *kelch13* mutations peaked at 141 in 2014, and unique validated/candidate mutations at 20 in 2012 (Figure S1). The majority of the increase in validated/candidate markers from 2000 - 2014 came from Southeast Asian populations (Figure S1). The increase in other propeller mutations in this time came mainly from Southeast Asia and also from West Africa.

By the time ACTs were introduced as first-line treatments in 2005, over half of the current WHO validated/candidate mutations had already been identified, predominantly in Southeast Asia (Figure 2). By this point, artemisinin derivatives were already widely used in the region and had presumably created an evolutionary pressure for these mutations to take hold. The beginnings of ART-R in Africa can be identified via the rapid increase of WHO validated/candidate marker diversity in Africa after 2010 (Figure 2). The cumulative count of WHO validated/candidate mutations in Africa was delayed relative to Southeast Asia and remained at three until 2010 (Figure 2). In Africa, each WHO validated/candidate mutation recorded before 2010 was only found in two samples. Such a small number leaves open the possibility that these mutations are artefacts, and suggests that ART-R did not begin to emerge in Africa until after 2010 (Figure 2). From 2010, the number of unique WHO validated/candidate mutations increased significantly to reach 16 of a possible 23 by 2018 (Figure 2). This means that ∼70% of WHO validated/candidate mutations have been detected in Africa at some point since 1980, though the vast majority arose after 2010, when the frequency of such mutations also began to increase, marking the beginning of the spread of ART-R in Africa.

**Figure 2.**
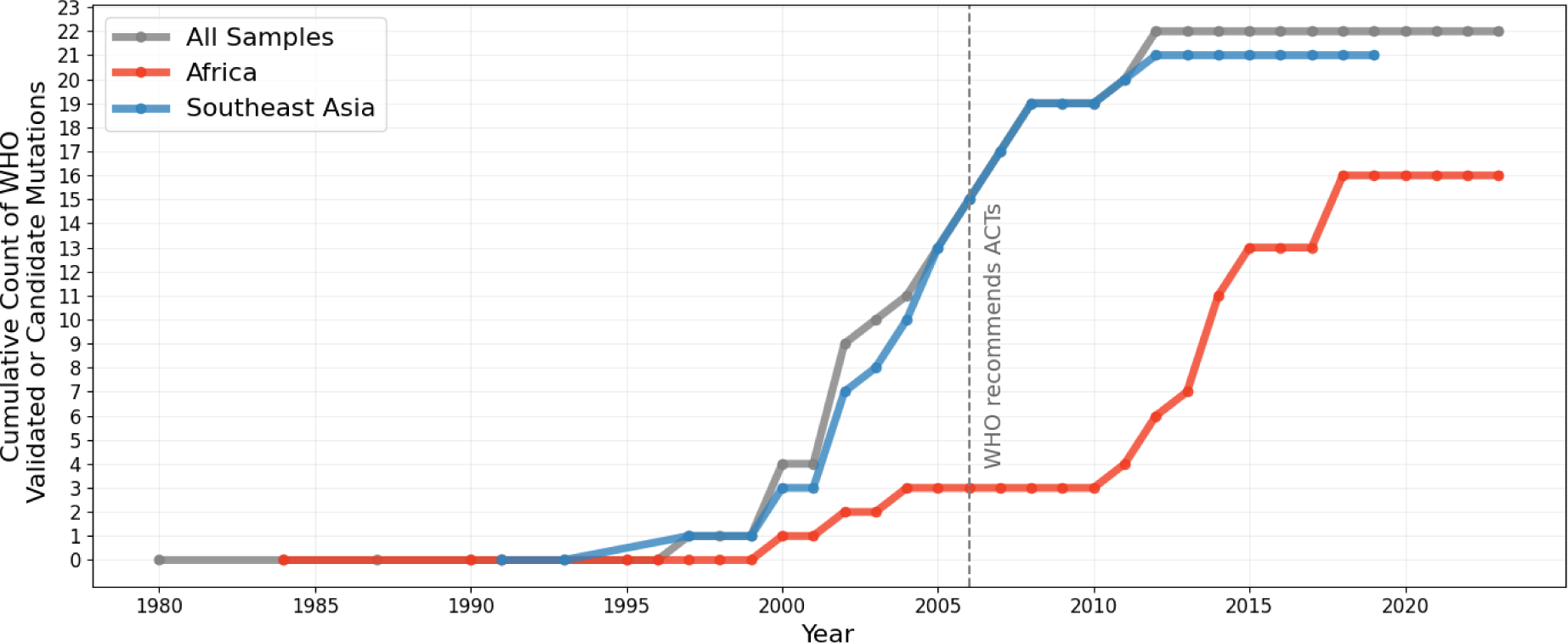
Cumulative count of unique WHO validated/candidate *kelch13* mutations over time across the whole dataset (grey), Southeast Asia (blue) and Africa (red).

In the past decade, sample sizes have declined, particularly in Southeast Asia, with most samples post-2019 coming from Africa. In contrast, the number of samples coming from West Africa dominated the dataset from 2014 - 2016, while the number of samples from Southeast Asia declined during this time (Fig 1C, Table S4). From 2020 - 2023, the average number of samples across all populations was 1,513, which was over ten times smaller than the number of samples collected in 2014 (Figure 1C, Table S4). From 2019, the drop in sample sizes may be explained by the lag in publication of more recently collected samples and possibly the difficulties in sample collection during the COVID-19 pandemic. However, these factors do not explain the sampling drop from 2014 - 2019. According to the 2023 world malaria report (WHO, 2023), total funding of malaria control in 2022 was US$3.7 billion below the estimated total required to meet global technical strategy targets for malaria control and elimination. It is possible that this inadequate funding has contributed to reduced sampling, however more evidence would be needed to establish this. There was no data from Southeast Asia in our dataset after 2019, but the proportion of mutant samples and variety of *kelch13* mutations was still highest in this region in 2019 (Figures 3 and 4).

**Figure 3.**
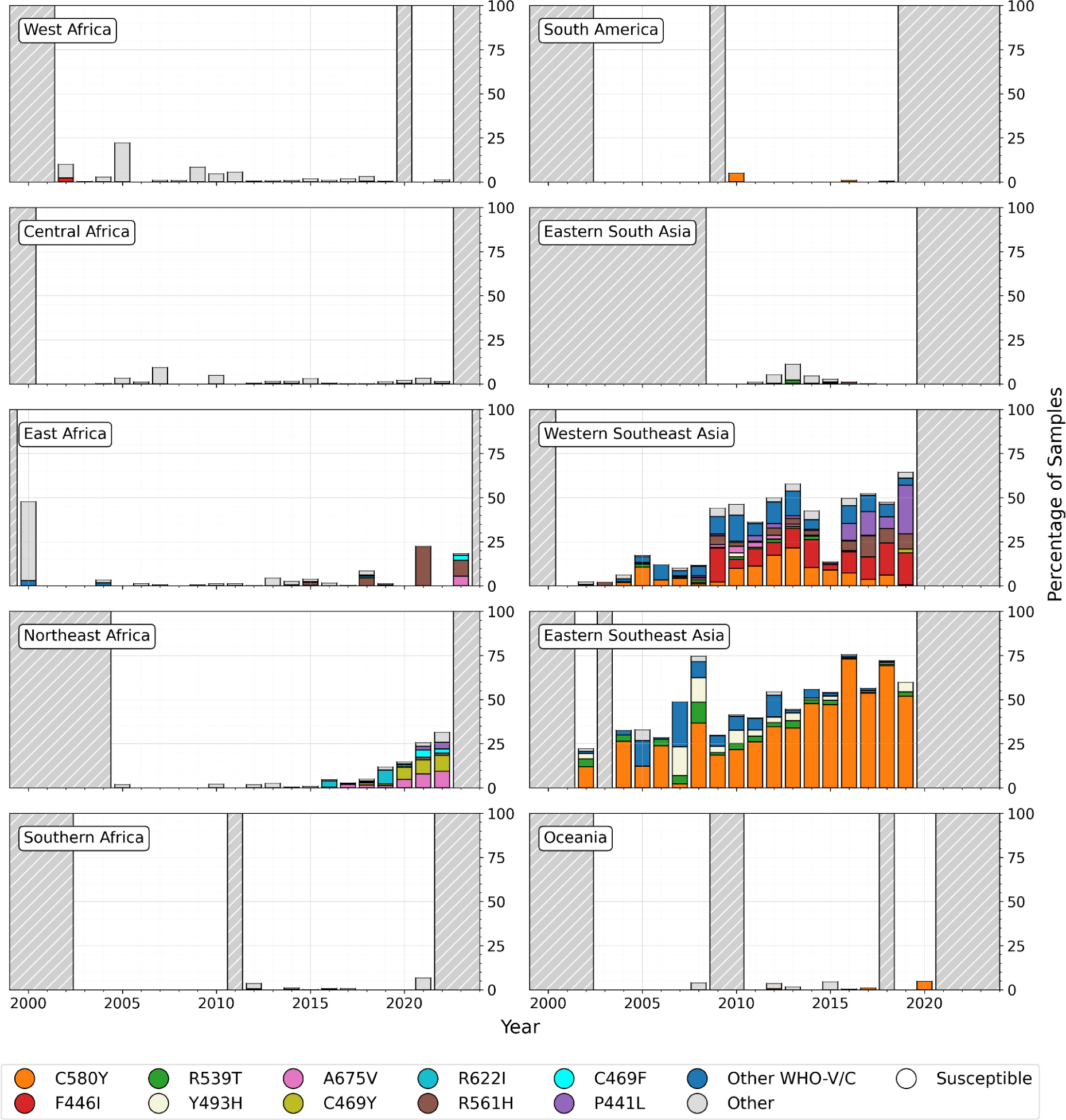
Proportions of *kelch13* mutations over time from 2000 - 2024 for ten populations, with African populations in the left-hand column. Years with fewer than 25 samples are shaded in grey with white stripes. Three populations (North Africa, Western Asia, Far-Eastern South Asia) are not displayed due to lack of data. Several of the most common WHO validated/candidate markers are highlighted. Other WHO validated/candidate markers are denoted by ‘Other WHO-V/C’. Mutations in the *kelch13* BTB/POZ or propeller domain without validated/candidate status are categorised as ‘Other’. The 3D7 reference *kelch13* sequence and the A578S mutation are known to not confer ART-R, so are categorised together as ‘Susceptible’. The high percentage of ‘Other’ mutations in East Africa in 2000 is due to our aggregation of samples from 1996-2003 to the median year of 2000 from a single study by de Laurent *et al*. (2018). Similarly, the percentages of ‘Other’ mutations in West Africa (2005) and Central Africa (2007) result from single studies (Ouattara *et al*., 2015; Taylor *et al*., 2015) where multiple different mutations were identified.

**Figure 4.**
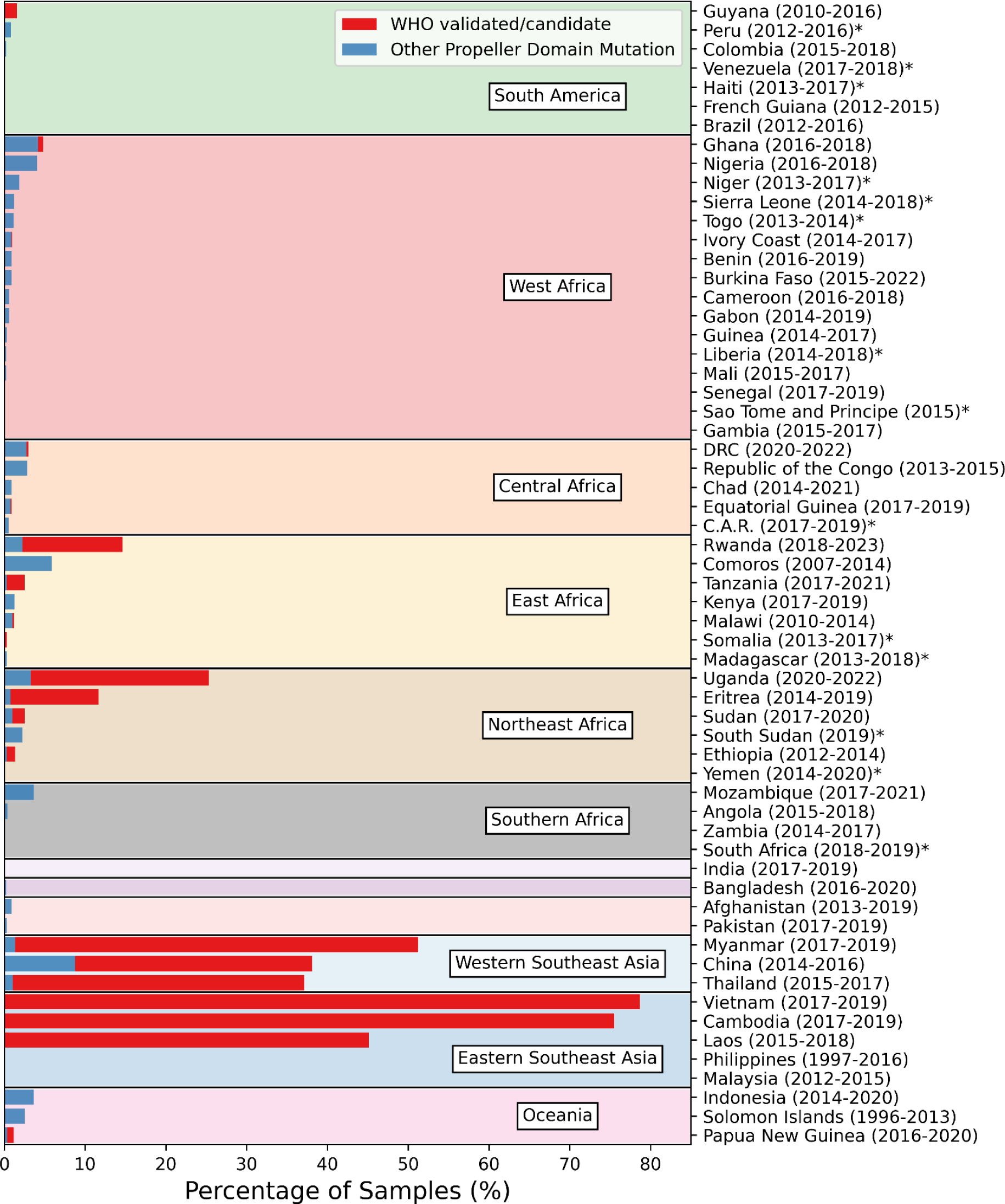
Proportion of samples from each country and geographical population with a WHO validated/candidate *kelch13* mutation (red) or other *kelch13* mutation in the BTB/POZ or propeller domain (blue). Bars show the weighted average percentage of samples from the three most recent observed years with at least 25 samples. Averages were weighted by the total number of samples observed between the years. Date ranges show the range of years from which the three most recent years with at least 25 samples were taken. Countries with fewer than 3 years with at least 25 samples are marked with an asterisk.

Since 2020, 83 unique *kelch13* propeller mutations have been reported globally, including eight WHO validated/candidate mutations. The majority of WHO validated/candidate mutations detected since 2020 occurred in East, Northeast, and Central Africa and no data was published after 2019 in Southeast Asia (Figure 3). The only population outside of Africa with at least 25 samples in a year from 2020 was Oceania, where WHO-validated C580Y was found at 5% in 2020 (Figure 3).

Northeast Africa had the highest diversity of unique validated/candidate mutations in Africa this decade (n = 7) and there has been a notable increase in the prevalence of several such markers in the region in this timeframe. The WHO-validated A675V doubled from 4.7% to 9.4% between 2020 - 2022. Another WHO-validated mutation, C469Y, showed a less pronounced increase, from 6.9% in 2020 to 9% in 2022. A mutation at the same position, WHO-candidate C469F, almost tripled in frequency, moving from 0.9% in 2020 to 2.5% by 2022. WHO-candidate P441L increased by nearly ten-fold this decade, though the final recorded frequency was relatively low, at 3.6% in 2022. Validated/candidate markers R539T and R622I were detected at low frequencies in 2020 (< 0.5%), and R561H was in 1.4% of Northeast African samples in 2022. In East Africa, multi-year sampling was rarer than in Northeast Africa. Three WHO validated/candidate mutations were detected in East Africa in 2023: C469F (2.8%), A675V (5.6%), and R561H (declined from 22.5% in 2021 to 8.9% in 2023). In Central Africa, validated/candidate markers P441L, R561H, and C469Y were each detected in single years, all below 0.5%.

The combined frequency of non-validated/candidate *kelch13* mutations also rose in Northeast Africa, from 2.5% in 2020 to 7.3% in 2022. While A578S (not associated with ART-R) contributed to these figures, several other propeller mutations were observed to be increasing in frequency in Northeast Africa: P441A, N490T, V637I, L713F, and A724P (though all frequencies were below 1%). The only non-validated/candidate mutation in East Africa was A578S, found at 0.9% in 2023. In Central Africa, V637I and S552C were found at a combined frequency of 0.67% in 2022. These data support the growing concerns that ART-R has begun to emerge in Africa. Several confirmed molecular markers of ART-R were either observed to have increased in frequency since 2020, or were observed at appreciable frequencies where longitudinal data were not available (Figure 3). The data also revealed increases in non validated/candidate *kelch13* propeller mutations. These newly arising mutations hold the potential to confer ART-R, and should be fast-tracked for assessment against the WHO validated/candidate marker criteria (Table S1).

### 2.4 Data Limitations

Compiling such a large dataset, which spans many locations and years, has allowed us to identify important trends in *kelch13* mutation sampling and emergence. It has also highlighted biases in the data. It is concerning that sampling effort has consistently reduced throughout the past decade (Figure 1C). As highlighted by Dhorda *et al*. (2024), the unfolding of resistance to antimalarial drugs must be closely monitored. Our analysis highlights a disparity between what should be happening for adequate surveillance of ART-R (routine, longitudinal sampling) and what has actually been occuring (opportunistic and reduced sampling). There were geographical biases in the dataset, with the majority of samples coming from Africa (58%) and Southeast Asia (29%). With these regions carrying most of the global malaria burden, this is understandable (WHO, 2023). Malaria eradication, which is the ultimate goal of the WHO (WHO, 2020), will require reliable insights from genomic surveillance from all regions in which malaria exists. Sampling bias can arise from various sources, including the variation in location or number of health facilities between regions (Mayor *et al*., 2023) and differing sampling regimes between national control programmes. In some cases, we see an over- and under-representation of some countries within the same population (Table S2). Moreover, while the majority of samples come from the continents most affected by malaria, there are key temporal imbalances within and between these continents. Global comparison of all populations was not possible after 2020 due to a lack of samples from Asia and South America (Figures 3 and 4). In Africa, sampling between regions has not been equal or consistent in the last ten years. For example, ten years ago in 2014, West Africa made up 52% of African samples but in 2019, this was just 5%. In Northeast Africa, these proportions were 9% in 2014 and then 25% in 2019. Especially in more recent years, since 2019, there was no year in which all of the regions most impacted by malaria (all except North Africa) were represented. Having systematic, longitudinal sampling in Africa, that is quickly made publicly available, will be crucial for providing robust insights into the evolution of malaria against control measures such as ACTs.

## 3. Spread of Artemisinin Partial Resistance

In the previous section we highlighted how ART-R has been found across the globe, but until recently, had been mostly localised in Southeast Asia. Concerningly, several WHO validated/candidate mutations have now emerged in parts of Africa, with potentially widespread consequences for human health. Here, we provide a more detailed case study for Southeast Asia and East/Northeast Africa, focusing on how resistance-associated *kelch13* markers emerged and spread, summarising their current frequencies, and highlighting key questions on their geographical distributions. We then discuss the *kelch13* substitutions observed in other regions of the globe, including West and Central Africa, South America, and Oceania.

### 3.1 Southeast Asia

#### 3.1.1 How did kelch13 mutants initially emerge and spread in Southeast Asia?

Artemisinin combination therapies have been used extensively across Southeast Asia since the mid-1990s, where they were quickly adopted by national governments seeking alternatives to drugs such as chloroquine, which had developed widespread resistance. Thailand, for example, adopted ACTs in the mid-1990s, shortly followed by Cambodia in 2000, Myanmar in 2002 and Vietnam in 2003 (Hassett and Roepe, 2019). Initially, there was optimism that ART-R might not occur, or at least that it would occur more slowly than resistance to previous drugs, due to the combination of an artemisinin derivative with a partner drug (Yeung *et al*., 2004; Pongtavornpinyo *et al*., 2008). However, this optimism faded in 2008 with the discovery of ART-R on the Thai-Cambodian border (Ashley *et al*., 2014; WHO, 2016; Tun *et al*., 2017). The proportion of samples with a validated/candidate marker had in fact already been high by 2006, before increasing in Thailand, Cambodia, and Myanmar, particularly between 2007-2016 (Figure 6) (Kagoro *et al*., 2022). ART-R associated *kelch13* mutations then began to emerge independently across several locations in Southeast Asia. Once ART-R had emerged, a hard selective sweep then drove a single multidrug resistant co-lineage, KEL1/PLA1, to dominance across the eastern Greater Mekong Subregion (Takala-Harrison *et al*., 2015; Imwong, Suwannasin, Kunasol, Sutawong, Mayxay, Rekol, Smithuis, Hlaing, Tun, Pluijm, *et al*., 2017; Amato *et al*., 2018; Hamilton *et al*., 2019; Imwong, Dhorda, Tun, *et al*., 2020). This lineage contained the C580Y *kelch13* mutation. Analysis of genomic regions flanking the *kelch13* gene suggested this migration occurred several times, with the C580Y mutation initially spreading eastward from Cambodia to Vietnam, and later, from Cambodia to Thailand and Laos around 2013-15 (Imwong *et al*., 2017, Takala-Harrison *et al*., 2015). Over the next 10-12 years, the C580Y mutation reached near-fixation frequencies in several of these countries (Figure 5). In contrast, in Myanmar, C580Y emerged and spread independently on a different parasite genetic background (Imwong, Suwannasin, Kunasol, Sutawong, Mayxay, Rekol, Smithuis, Hlaing, Tun, van der Pluijm, *et al*., 2017). This pattern of identical *kelch13* mutations arising independently, alongside migration events between countries would become a repeated occurrence across Southeast Asia, with several of the most common *kelch13* mutations, including C580Y, spreading via both mechanisms (Takala-Harrison *et al*., 2015; Imwong *et al*., 2017; Hamilton *et al*., 2019; Imwong *et al*., 2020).

**Figure 5.**
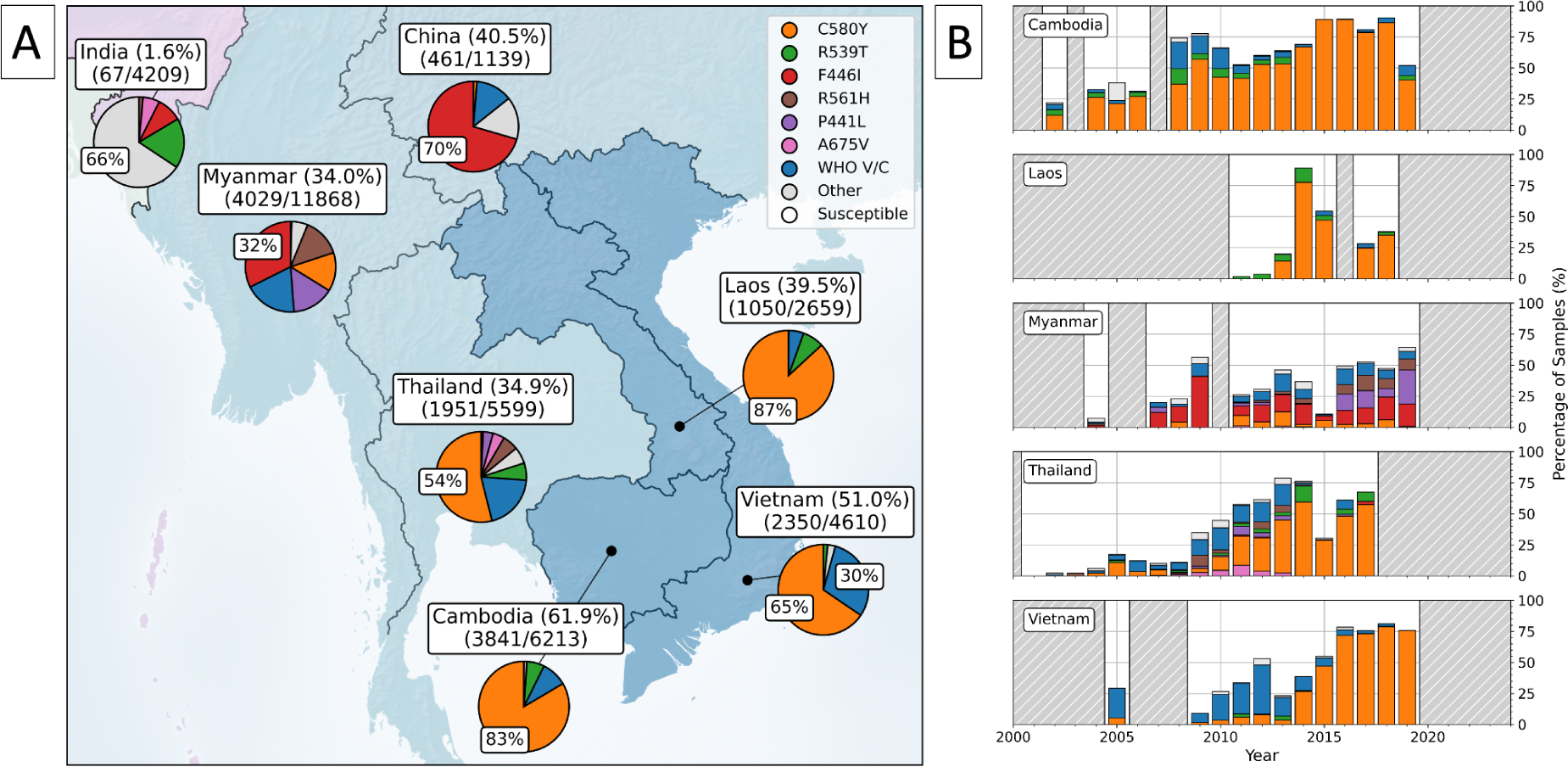
Panel A shows the distribution of *kelch13* propeller mutations across Southeast Asia. The percentage of samples with any *kelch13* propeller mutation are included in the country labels, above the number of samples with any *kelch13* propeller mutation and the total number of samples collected for each country, respectively. Among only the samples with any observed *kelch13* propeller mutation, pie charts show the proportions with markers of interest, where proportions above 25% are labelled accordingly. ‘Other’ denotes low frequency mutations in the propeller domain which are not WHO validated/candidate markers, aggregated into a single category. Samples with the 3D7 reference sequence for *kelch13* or A578S are denoted as ‘Susceptible’ (WHO, 2022). Panel B shows all samples collected over time for each country as a stacked bar chart, where the proportion of samples with each marker is coloured. Years with fewer than 25 samples are highlighted with grey dashed lines.

**Figure 6.**
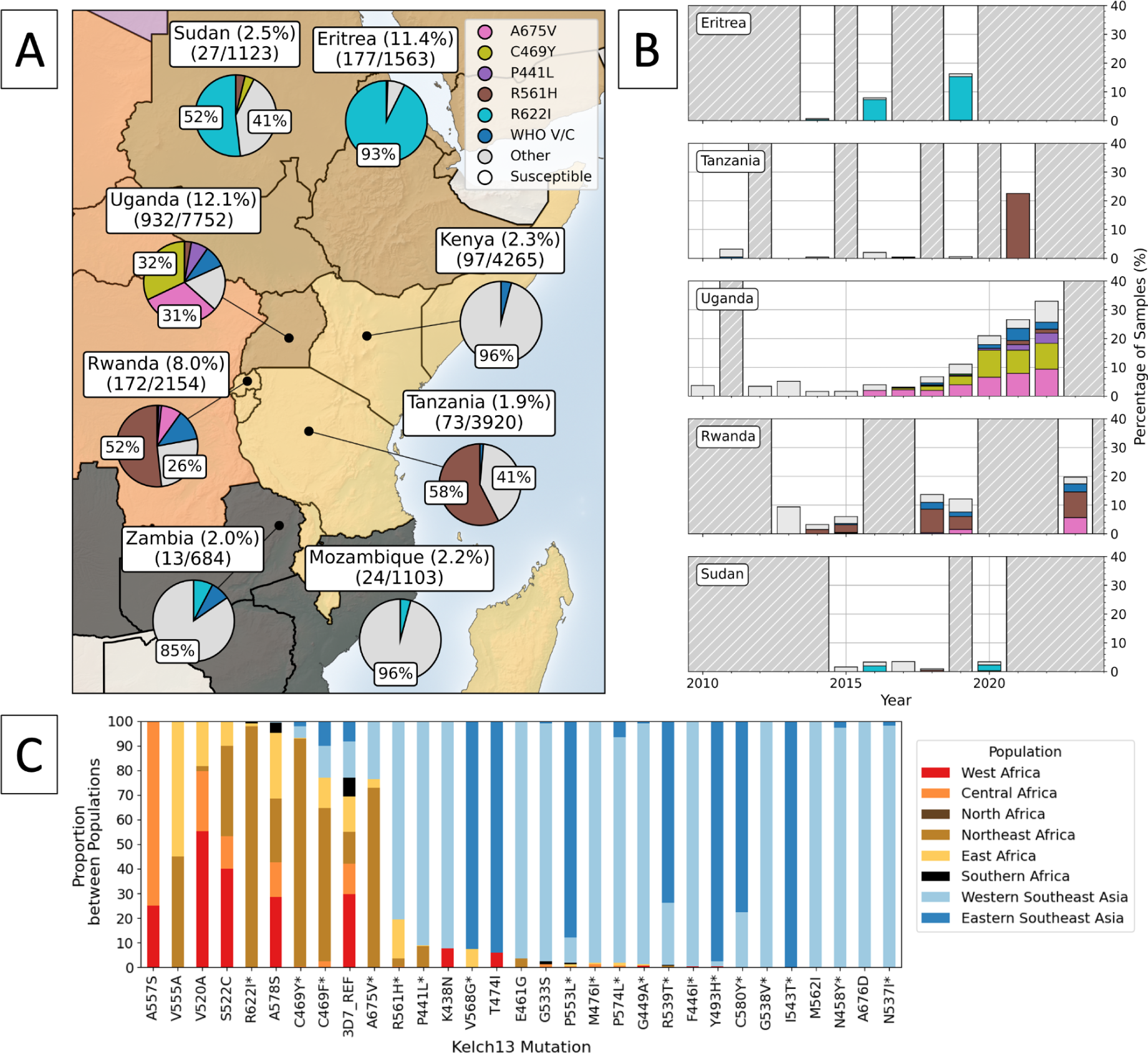
Panel A shows the distribution of *kelch13* propeller mutations across East and Northeast Africa. The percentage of samples with any *kelch13* propeller mutation are included in the country labels, above the number of samples with any *kelch13* propeller mutation and the total number of samples collected for each country, respectively. Among only the samples with any observed *kelch13* propeller mutation, pie charts show the proportions with markers of interest, where proportions above 25% are labelled accordingly. ‘Other’ denotes low frequency mutations in the propeller domain which are not WHO validated/candidate markers, pooled into a single category. Samples with the 3D7 reference sequence for *kelch13* or A578S are denoted as ‘Susceptible’ (WHO, 2022). Panel B shows all samples collected over time for each country as a stacked bar chart, where the proportion of samples with each marker is coloured. Years with fewer than 25 samples are highlighted with grey dashed lines. Notably, Uganda and Rwanda have shown a consistent increase in diversity among *kelch13* mutations between 2016-2023. Panel C shows all mutations with at least 25 samples across Africa and Southeast Asia on the axis. Coloured bars show which subpopulation samples came from across Africa and Southeast Asia. WHO validated or candidate mutations are suffixed with an asterisk.

Genomic epidemiology played a fundamental role in identifying these patterns of emergence and migration in Southeast Asia. While it was initially thought there had been a single emergence of ART-R in the region, comparison of parasite genetic backgrounds revealed it had in fact emerged independently across several countries (Amato *et al*., 2018; Verity *et al*., 2020; Neafsey, Taylor and MacInnis, 2021). This enabled policy makers to focus on slowing local resistance through improved surveillance and access to malaria prevention, rather than focusing solely on preventing parasite migration across borders (WHO, 2015b; Manzoni *et al*., 2024). Data from surveillance efforts, including molecular surveillance, then became a cornerstone for developing mitigation strategies in the region. One such strategy, the ‘WHO Strategy for Malaria Elimination in the Greater Mekong Subregion’ (2015–2030), leveraged this evidence to achieve significant reductions in malaria burden of 77% (WHO, 2015b; Manzoni *et al*., 2024). This underscores the importance of accurate and consistent molecular surveillance data in formulating effective public health policies and interventions.

#### 3.1.2 What is the current distribution and frequency of kelch13 mutations across Southeast Asia?

The proportion of samples with a WHO validated or candidate *kelch13* substitution remained high as of 2019 - the most recently sampled year in Southeast Asia in our dataset. Based on the latest data, between 37.9 - 75.8% of samples had a validated/candidate mutation in most Southeast Asian countries (Figure 5). For example, the proportion of samples with a WHO validated/candidate mutation in Vietnam (2019) and Thailand (2017) was 75.8% and 67.5% respectively (Table S5). Similarly, in Cambodia, these frequencies fluctuated between 80-90% from 2016 - 2018. In 2019, 52.1% of Cambodian samples had a WHO validated/candidate mutation, however, data for this year were collected from three sites in only one study, compared to between six and eleven sites from several studies in 2016 - 2018 (Peto *et al*., 2022). Laos showed a slightly lower frequency of 37.9% (2018), though exhibited significant year-to-year variability, with a peak of 89% in 2014 (based on 111 samples). In contrast, Myanmar showed both a lower frequency and slower increase in the proportion of samples with a *kelch13* mutation between 2004 and 2019 (Table S5), though frequencies were still high as of the most recent year (51.5%). Between 1993 and 2019, there were 117 unique *kelch13* mutations present across Southeast Asia, though only 22 of these were observed in more than 25 samples (Table 1). While the C580Y mutation, a WHO-validated marker, was initially only one of several markers to emerge, it later rose to high frequency in many countries, such as Thailand, Laos, Cambodia, and Vietnam.

**Table 1.** Most common *kelch13* mutations across Southeast Asia, where mutations occur in at least 25 samples across the two populations (western and eastern Southeast Asia). Countries represented in this summary are Myanmar, China, Thailand, Cambodia, Laos and Vietnam. Total refers to the total number of samples from these countries with each marker.

| Marker | WHO status | Total | Proportion (%) |
| --- | --- | --- | --- |
| 3D7 reference sequence | - | 18,958 | 58.08 |
| C580Y | validated | 7,271 | 22.27 |
| F446I | validated | 1,644 | 5.04 |
| P441L | candidate | 672 | 2.06 |
| R561H | validated | 666 | 2.04 |
| R539T | validated | 478 | 1.46 |
| Y493H | validated | 437 | 1.34 |
| I543T | validated | 391 | 1.20 |
| G449A | candidate | 355 | 1.09 |
| P574L | validated | 292 | 0.89 |
| P553L | validated | 226 | 0.69 |
| M476I | validated | 173 | 0.53 |
| N458Y | validated | 140 | 0.43 |
| G538V | candidate | 126 | 0.39 |
| A675V | validated | 95 | 0.29 |
| G533S | no status | 86 | 0.26 |
| N537I | candidate | 49 | 0.15 |
| A676D | no status | 44 | 0.13 |
| V568G | candidate | 38 | 0.12 |
| M562I | no status | 35 | 0.11 |
| T474I | no status | 31 | 0.09 |
| C469F | candidate | 30 | 0.09 |
| E461G | no status | 27 | 0.08 |

As of 2019, C580Y remains dominant, with the majority of samples with any *kelch13* mutation showing this marker. For instance, 83% of all samples from Cambodia with a *kelch13* mutation had C580Y. A similar trend was observed in Thailand (54%) and Vietnam (65%). While other *kelch13* substitutions were circulating in these countries, such as R539T and R561H, these were at much lower frequencies. In Thailand, C580Y had a smaller majority than Cambodia, Vietnam, or Laos and a larger diversity of other markers. This is most likely due to the malaria-free corridor which separates parasite populations in east and west Thailand. Eastern Thailand tends to have higher proportions of C580Y while western Thailand more closely reflects the diversity in bordering Myanmar (Kobasa *et al*., 2018). In Myanmar, several unique *kelch13* mutations were circulating at moderate frequencies in 2019, with no single substitution reaching fixation. C580Y is not the dominant mutation in Myanmar, with mutations such as F446I, R561H, and P441L circulating, particularly since 2016, and at higher frequencies than neighbouring Thailand (Bonnington *et al*., 2017). This pattern of several circulating markers has remained consistent between 2004-2019, whereas countries such as Thailand and Laos saw C580Y overtake others in frequency during the same time period. In total, of the 22 most common *kelch13* mutations, 8 of these were unique to Myanmar and China within Southeast Asia, including: E461G, A675V, M562I, F446I, A676D, G538V, R561H, P441L (Table 1). A further 6 mutations were more common in these countries than eastern Southeast Asia: M476I, G449A, G533S, N537I, N458Y, and P574L. Only three mutations were unique to the Eastern part of Southeast Asia: I543T, V568G, and T474I, though a further four were more common than in Myanmar/China (R539T, C580Y, P553L, and Y493H). However all of these were at low frequencies in these countries.

The high frequency of *kelch13* mutations in Southeast Asia has translated into increased negative clinical outcomes, as evidenced by delayed clearance rates above 10% in TES (Table S6). We obtained the results of studies which monitor the therapeutic efficacy of common ACTs from Malaria Threats Map (https://apps.who.int/malaria/maps/threats/ - WHO, 2024) (Figure S2). Out of 94 studies across Southeast Asia, 52 (55%) reported delayed clearance rates (>10%) for dihydroartemisinin-piperaquine (DHA-PPQ) treatment, with 18 (19%) reporting treatment failure rates over 10%. Artesunate-mefloquine (AS-MQ) paints a similar picture: of 38 studies, 21 (55%) reported delayed clearance rates, with 2 (5%) reporting treatment failure rates over 10%. In contrast, artemether-lumefantrine (AL) has shown slightly better outcomes, with only 15 of 131 studies (11%) reporting delayed clearance rates of >10%, and four (3%) reporting treatment failure rates of >10%. This lower delayed clearance rate for AL may be attributed to the fact that AL has only been a first-line treatment in Laos and Myanmar, where ART-R is lower (WHO, 2015b, 2021; Jacob *et al*., 2021; WHO, 2023). The high frequency of delayed clearance and treatment failures indicate the growing challenge in malaria treatment using ACTs in Southeast Asia.

These findings underscore the need for ongoing genomic surveillance to establish the composition and frequencies of circulating parasite populations in Southeast Asia. However, data from the region are also limited by the lack of systematic, longitudinal sampling, without which it is difficult to properly estimate the frequencies of resistance within a given area (Mayor *et al*., 2023). For example, in our data, no information was available for Southeast Asian countries after 2019. Additionally, many countries showed considerable between-year variation in the proportion of samples with *kelch13* mutations, which may be due to a disproportionately large number of samples from a single study or region and inconsistent sampling of regions. One major challenge to this call for more longitudinal sampling is that it becomes more challenging as case numbers are driven down, as is the case in Southeast Asia. While systematic genomic surveillance is still key to properly interpreting the temporal frequency trends of *kelch13* mutations, going forwards, additional strategies may also be required to account for lower case numbers in regions near to elimination. These could include enhanced collaborations and data sharing to pool data, and compensatory statistical techniques like time-aggregated sampling.

#### 3.1.3 Did fitness costs affect the spread of kelch13 mutations across Southeast Asia?

Interestingly, differences in the clearance rate and fitness effects of different *kelch13* mutations are correlated with their geographic distribution across Southeast Asia (Straimer *et al*., 2015; Stokes *et al*., 2021). In our dataset, we saw mutations associated with a longer parasite clearance time and larger fitness costs predominantly in Eastern Southeast Asia, mutations with moderate levels across the whole region, and mutations with lower ART-R effects and fitness costs restricted to Western Southeast Asia (Figure 5).

For example, the R539T mutation has a larger effect in ring-stage survival assays (RSA) than C580Y, but it also generates a larger fitness cost *in vitro* (Ashley *et al*., 2014; Straimer *et al*., 2015; Stokes *et al*., 2021). In our dataset, 98% of samples with R539T were from Eastern Southeast Asia, including eastern locations in Thailand. C580Y is a more prevalent generally than R539T across Eastern Thailand, Laos, and Cambodia and is also found in more samples from Western Southeast Asia, in Myanmar (Anderson *et al*., 2017; Imwong, van der Pluijm, *et al*., 2017). C580Y is less common than F446I in Myanmar and Western Thailand (Figure 5). F446I has been shown to confer a lower degree of ART-R but also a lower fitness cost in competition assays than C580Y (Ashley *et al*., 2014; Stokes *et al*., 2021). Compared to other locations in Southeast Asia, there is known to be increased transmission, lower drug pressure, and higher competition between parasites in Myanmar, which makes up the majority of cases from Western Southeast Asia. The lower fitness cost of the F446I mutation may provide a selective advantage over C580Y in these areas (Imwong *et al*., 2017b, 2020). Similarly, R539T may have been able to emerge in Eastern Southeast Asia, because competition is lower than Myanmar, while the moderate ART-R and fitness cost of C580Y may have enabled it to spread more widely in Southeast Asia.

These effects may have resulted in several mutations with lower fitness costs being more strongly selected for than mutations which cause greater ART-R, potentially explaining the relatively stable frequencies of several mutations observed there between 2004-2019. In a further example, CRISPR/Cas9-edited parasites with R561H show similar survival rates to parasites with engineered C580Y mutations under treatment, but a decreased fitness cost *in vitro* (Nair *et al*., 2018). If increased survival rates and lower fitness costs do co-occur, it may mean R561H has potential to spread widely among populations, which is concerning as it has increased in frequency in Myanmar since 2016 (Imwong *et al*., 2017a; Hamilton *et al*., 2019).

Notably, individual *kelch13* substitutions have been shown to not only have different phenotypic and fitness effects, but that these effects can differ depending on the genetic background of the parasite in question (Siddiqui, Liang and Cui, 2021; Stokes *et al*., 2021; Stokes, Ward and Fidock, 2022). For example, gene-editing studies have demonstrated that after introducing *kelch13* mutations into parasites of Southeast Asian origin, they exerted a lower fitness cost and a greater effect on RSA survival than parasites from Africa (Stokes *et al*., 2021; Stokes, Ward and Fidock, 2022). There is also evidence of ‘genetic backgrounds’ which are associated with *kelch13* mutations, and tend to be more common in parasites from areas in Southeast Asia where ART-R has emerged and spread extensively (Miotto *et al*., 2015; Cerqueira *et al*., 2017; Uwimana *et al*., 2021). For example, genomic changes have been identified which may compensate for the fitness costs of the *kelch13* mutations, such as by altering nutrient permeable channels in the parasite vacuole membrane, and allowing uptake of important amino acids. These backgrounds are more common in Southeast Asia, which could be one possible reason why ART-R emerged and spread in this region so quickly, alongside early and widespread ACT use (Mesén-Ramírez *et al*., 2021). Taken together, these effects could mean resistance mutations are more likely to appear (and therefore spread) on certain genetic backgrounds, or in combination with important genetic changes (Miotto *et al*., 2015; Cerqueira *et al*., 2017; Stokes *et al*., 2021).

As novel *kelch13* mutations emerge and spread, understanding their fitness costs across different geographical regions and in relation to parasite genetic backgrounds will be crucial. This understanding will require assessing fitness costs through in-vitro assays and integrating these findings with genomic surveillance to reveal broader patterns. Genomic surveillance may aid in generating models which can predict the spread of resistance given the frequency of certain genetic backgrounds. These patterns also highlight the advantages of whole genome sequencing over targeted amplicon sequencing, as this additional information can be used to inform models using the genetic background of resistant parasites.

### 3.2 East and Northeast Africa

#### 3.2.1 How did kelch13 mutants initially emerge and spread in East and Northeast Africa?

ACTs have been used extensively across Africa, particularly after 2006 (Tables S6 and S7). By 2008, almost all African countries had implemented ACTs as the primary treatment for malaria, meaning the emergence and spread of ART-R in Southeast Asia around 2008 occurred in tandem with a growing reliance on ACTs in Africa (WHO, 2009). This led to fears that similar mutations could not only arise in Africa, but would result in a wave of resistant parasites, with dramatic consequences for health in the region. In the last five years, those fears are beginning to be realised, with reports of WHO validated/candidate *kelch13* mutations in several countries across East and Northeast Africa. This started with the identification of the R561H mutation in Rwanda in 2019. This mutation was shown to have been circulating in the region since 2013-2015, before increasing to ∼13% in 2019, with demonstrated effects on parasite clearance rate (Uwimana *et al*., 2020, 2021). This was then mirrored by the dramatic increase in both A675V and C469Y mutations in Uganda between 2015-2019, which were also associated with delayed parasite clearance in the country (Balikagala *et al*., 2021). Given the extent of ACT use in Africa over the past two decades, it is unclear why ART-R has only emerged now, around 10-15 years behind Southeast Asia, though it is most likely due to the slower initial rollout of ACTs in Africa, combined with differences in case-by-case drug exposure (Hassett and Roepe, 2019). Other factors, such as higher immunity, intra-host competition, or effective population sizes in Africa may also have slowed ART (see Section 4.1.3) (Wasakul *et al*., 2023).

After detecting ART-R associated *kelch13* mutations in Africa, an initial concern was that these mutations had been introduced via gene flow from Southeast Asian parasites, as was the case with resistance to chloroquine and sulfadoxine–pyrimethamine (Wellems and Plowe, 2001; Gregson and Plowe, 2005). However, evidence from genomic surveillance suggests that the *kelch13* mutations observed so far in Africa have emerged independently. For example, parasites in Uganda determined to be ART-R with the ring-stage survival assay, including one with A675V, are genetically more similar to African parasites than those from Southeast Asia, suggesting an independent African emergence (Ikeda *et al*. 2018). Similarly, analysis of *kelch13* flanking microsatellites and genome-wide single-nucleotide polymorphisms suggest Ugandan parasites with C469F, C469Y, and A675V form a clade distinct from Southeast Asian parasites, implying a single evolutionary origin and clonal expansion within Africa (Conrad *et al*. 2023). A further example is A675V, C469Y and R561H mutants from Uganda, Rwanda and Tanzania, which have different *kelch13* flanking haplotypes to parasites with these mutations from Southeast Asia, again pointing to a single evolutionary origin which arose independently in Africa (Uwimana *et al*. 2020, Balikagala *et al*., 2021, Ishengoma *et al*., 2024). Together, these analyses show clear evidence of independent emergence of *kelch13* markers in Africa, suggesting local conditions are prompting the emergence and selection of ART-R. Nevertheless, ongoing genomic surveillance will be needed to identify migratory pathways between continents should they occur, both for *kelch13* mutations and their genetic backgrounds.

#### 3.2.2 What is the current distribution and frequency of kelch13 mutations in East and Northeast Africa?

As of 2024, WHO validated/candidate mutations have now been identified in several countries across East and Northeast Africa, including Uganda, Rwanda, Sudan, Eritrea, Tanzania, Kenya and Ethiopia (Fola *et al*., 2023 - data external to our dataset), plus several other neighbouring countries at low frequencies (Figure 6). Concerningly, the frequency of these mutations has increased dramatically since 2018, particularly in Uganda, Rwanda, Tanzania and Eritrea. From our data for 2022, 26% of samples from Uganda had a WHO validated/candidate mutation, an increase from ∼1% in 2013. Similarly, 17% and 15% of samples from Rwanda and Eritrea had a WHO validated/candidate mutation in 2023 and 2019, respectively. Tanzania also had a high proportion of parasites with a WHO validated/candidate mutation in 2021 (23%), increasing from ∼0.1-0.2% between 2017 and 2019. However, Tanzanian samples from 2021 were all from the Kagera Region, close to the borders with Uganda and Rwanda, which also have high levels of ART-R. Earlier Tanzanian samples were collected further from Uganda, so the increase in Tanzania could be partly due to sampling bias. Sudan shows a small proportion of parasites with a WHO validated/candidate mutation, reaching 2.2% in 2020. Similarly, neighbouring countries of Kenya, DRC, Zambia and Mozambique each showed a small proportion of samples with an WHO validated/candidate mutation, though the total frequencies of samples with any propeller domain mutation in these countries remained ∼4% or less. Concerningly, these trends are strikingly similar to those observed in Southeast Asia around 2008, where the initial emergence of WHO validated/candidate mutations was followed by exponential increases in the frequency of resistant parasites over a decade. Tracking these changes in ART-R is crucial, as increased ART-R increases selective pressure for resistance to partner drugs, raising the likelihood of overall treatment failure. Given there are limited alternatives to ACT use, if this pattern continues in East and Northeast Africa, this could have disastrous consequences for malaria control in the region.

Alongside an increase in WHO validated/candidate mutations, many East and Northeast African countries also had an increase in frequency of propeller domain *kelch13* mutations which are not assigned WHO validated/candidate status, including Kenya, Rwanda, Tanzania, Eritrea, South Sudan, Sudan, and Uganda. The phenotypic effects of these mutations are not clear, but in some cases *in vitro* evidence suggests they could have phenotypic effects on artemisinin clearance rates (Mukherjee *et al*., 2017; WWARN K13 Genotype-Phenotype Study Group, 2019). Tracking the changes in frequency of these mutations is important, as they may in fact provide some degree of ART-R.

As of 2024, we have not seen any single WHO validated/candidate mutation rise to prominence in East or Northeast Africa, though this may occur in coming years - as with C580Y in Southeast Asia. In the past five years, several WHO validated/candidate markers have circulated simultaneously, with individual mutations mostly localised to specific countries within the region, similar to patterns seen in the early period of ART-R in Southeast Asia (Figure 6 and Table 2). For example, the most common WHO validated/candidate mutation, A675V, has risen to high frequencies in Uganda and Rwanda (9.4% in 2022 and 5.6% in 2023, respectively), but has not been observed in neighbouring countries. Similarly, Rwanda and Tanzania have a high frequency of parasites with the R561H mutation (8.9% in 2023 and 22.5% in 2021), but this mutation is uncommon elsewhere. A third example is R622I, which was the most common mutation in Eritrea (15.2% in 2019) and Sudan (2% in 2020), but rare outside Northeast Africa. Several of these mutations have increased in frequency over the past 5 years, with A675V, C459Y and P441L all increasing in frequency in Uganda since 2018. A similar situation appears to have occurred recently in Rwanda, where three validated/candidate mutations increased in frequency between 2019 and 2023; A675V from 1.5% to 5.6%, C469F from 1.5% to 2.8%, and R561H from 4.5% to 8.9%. Some countries also showed high diversity in the WHO validated/candidate mutations circulating, such as Uganda, which had six unique WHO validated/candidate mutations: P441L, C469F, C469Y, R539T, R561H, and A675V, likely due to the increased sampling there since 2019 (Conrad *et al*., 2023). The emergence and spread of different *kelch13* mutations in separate regions of East and Northeast Africa suggests they are so far emerging independently in response to local conditions, rather than migrating between areas (Abera *et al*., 2021; Assefa, Fola and Tasew, 2024).

**Table 2.**
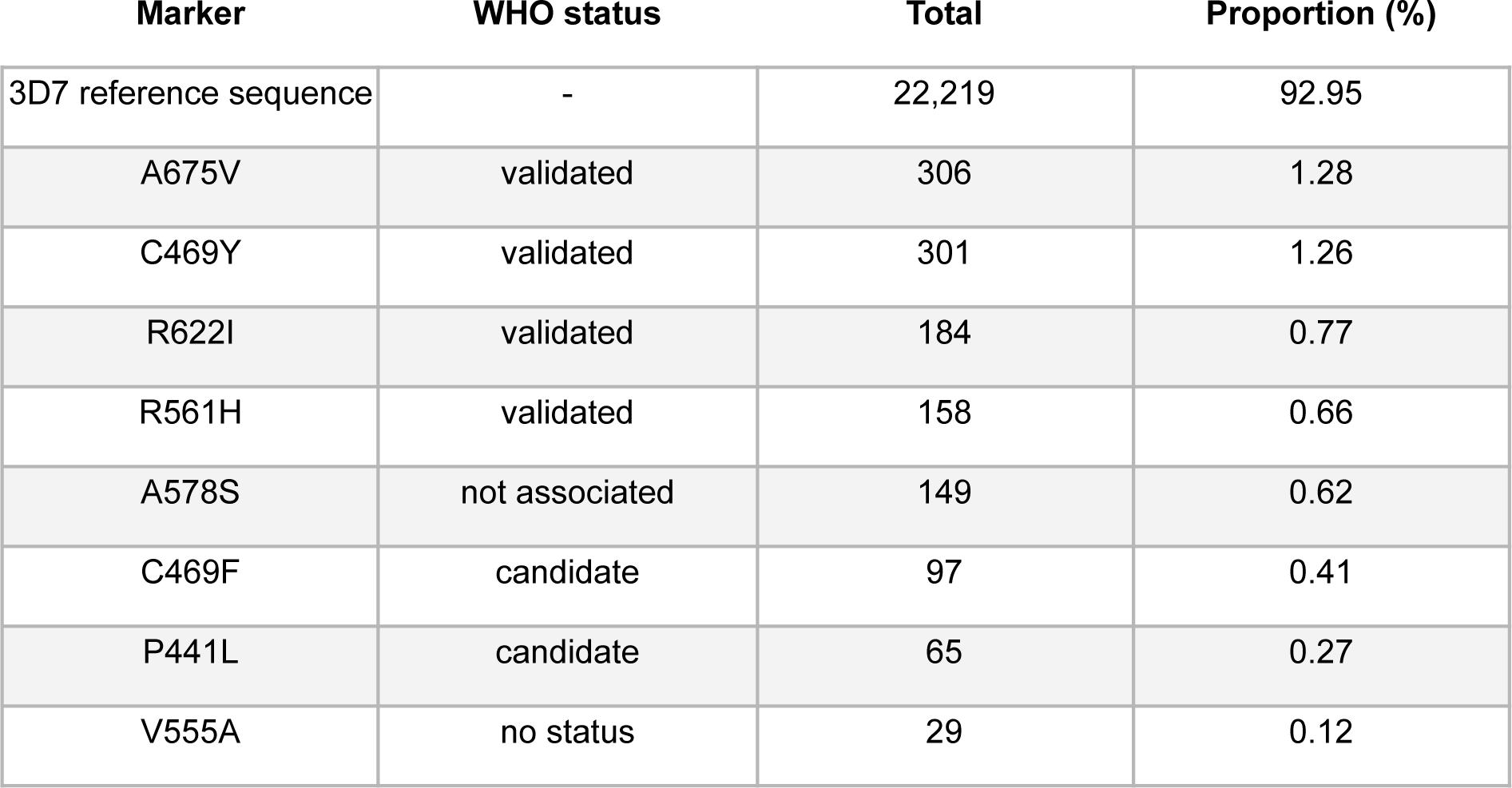
Most common *kelch13* mutations across East and Northeast Africa, where mutations occur in at least 25 samples. Countries represented in these populations are Burundi, Comoros, Kenya, Madagascar, Malawi, Rwanda, Somalia, Tanzania, Eritrea, Ethiopia, Saudi Arabia, South Sudan, Sudan, Uganda, Yemen. Note A578S is not associated with ART-R (WHO, 2022). Total refers to the total number of samples from these countries with each marker.

| Marker | WHO status | Total | Proportion (%) |
| --- | --- | --- | --- |
| 3D7 reference sequence | - | 22,219 | 92.95 |
| A675V | validated | 306 | 1.28 |
| C469Y | validated | 301 | 1.26 |
| R622I | validated | 184 | 0.77 |
| R561H | validated | 158 | 0.66 |
| A578S | not associated | 149 | 0.62 |
| C469F | candidate | 97 | 0.41 |
| P441L | candidate | 65 | 0.27 |
| V555A | no status | 29 | 0.12 |

Interestingly, the WHO validated/candidate *kelch13* mutations which have arisen in East and Northeast Africa are different to those found in Southeast Asia (Figure 6C). For example, the most common *kelch13* mutations in Southeast Asia are C580Y and F446I, but these are uncommon in Africa, with C580Y only observed in 7 samples across the entire continent. Other common substitutions in Southeast Asia were also rare in Africa, including R539T, P574L, F446I, and G449A, all with fewer than 5 samples. Similarly, of the most common WHO validated/candidate markers in Africa (A675V, C469Y, R622I, R561H, and C469F), only R561H was found regularly in Southeast Asia. This could be due to differences in ACT use between continents. For example, artemether-lumefantrine is the most common treatment for malaria in Africa, but in Southeast Asia, dihydroartemisinin-piperaquine and artesunate-mefloquine are used more often (Tun *et al*., 2018; WHO, 2022) (Table S7). Theoretically, these differences could promote different *kelch13* mutations, because their partner drugs are associated with selection for different genetic backgrounds with differing resistance mutations. For example, R539T has been associated with mefloquine-resistant parasites, whereas C580Y has been associated with piperaquine-resistant parasites, because changes in *mdr1* and *plasmepsin2/3* respectively alter the fitness costs of *kelch13* mutations in the presence of these different drugs (Parobek *et al*., 2017). It is also possible that differences in the phenotypic effects of the substitutions, alongside variation in transmission rates, fitness costs or genetic backgrounds in Africa have selected for different *kelch13* mutations relative to Southeast Asia. However, the importance of this difference in mutations, and the factors which have given rise to them remains mostly speculative, particularly with regard to how this would affect mitigation strategies going forward.

Genomic surveillance had a vital role in identifying WHO validated/candidate mutations circulating at low frequencies in Africa, acting as an early-warning system for ART-R with improved resolution over methods such as TES (Assefa, Fola and Tasew, 2024). For example, samples with validated/candidate markers were identified by genomic surveillance in Eritrea in 2014, Tanzania in 2017, and Rwanda in 2014, whereas changes in therapeutic efficacy did not occur until 2019, 2022, and 2018 respectively (Figure S2). The warning from genomic surveillance can provide public health bodies with additional time to prepare strategies and organise resources to combat the spread of ART-R (WHO, 2022a; Assefa, Fola and Tasew, 2024). It is critical to continue genomic surveillance in Africa, as it could provide policy makers further opportunities to prepare for coming waves of ART-R.

#### 3.2.3 Why is ART-R centred in Eastern and Northeastern Africa?

It is unclear why WHO validated/candidate mutations have arisen in Northeast and East Africa first, or why we have not seen a wider spread of these mutations across Africa, despite extensive ACT use. Resistance to chloroquine and SP also arose first in East Africa before spreading through the continent. However, in contrast to ART-R, the origins of chloroquine and SP resistance are thought to have been gene flow from Southeast Asia, while ART-R appears to have emerged independently (Curtis, Duraisingh and Warhurst, 1998; Mberu *et al*., 2000; Wellems and Plowe, 2001; Conrad *et al*., 2023). If not introduced to East and Northeast Africa from elsewhere, ART-R could possibly have arisen here first due to higher drug pressure in the region. For example, household survey data from 2003-2015 suggests ACT use among symptomatic children was significantly higher in Eastern Africa compared to Central or West Africa, particularly around 2015 (Bennett *et al*., 2017). During this time in Uganda, 70.2% of symptomatic children received an ACT, compared to an average of 19.7% across the continent (Bennett *et al*., 2017). Inappropriate use of artemisinin monotherapies has also been observed in some areas of Uganda, with inadequate dosing of artemether-lumefantrine observed in ∼12% of children and ∼16% of adults in some districts (Wang *et al*., 2018). Anecdotal reports also suggest this issue occurs more widely across Africa. Another possibility is unstable transmission rates of parasites in East and Northeast Africa could promote the development of ART-R, because when there is a switch from low to high malaria incidence in an area, host immunity is temporarily weakened and drugs need to be used more frequently (Rosenthal *et al*., 2024). This is relevant in East and Northeast Africa, as there are many highland regions which are known to be centres of both elevated and unstable transmission (Rodó *et al*., 2021). This in particular may have been a factor in northern Uganda, where the discontinuation of an effective indoor residual spraying program could have contributed to a sudden resurgence in malaria cases and an increase in drug pressure (Conrad *et al*., 2023). Indeed, Uganda accounted for 5% of all global malaria cases in 2022, with higher transmission rates relative to many other African countries (only Nigeria and the DRC had more cases in that year) (WHO, 2023).

#### 3.2.4 Could fitness costs or genetic backgrounds slow the emergence of ART-R in Africa?

Although fitness costs of *kelch13* mutations are interesting conceptually, what potential do they have to slow the emergence of ART-R in Africa? Despite WHO validated/candidate *kelch13* mutations having arisen several times in East and Northeast Africa, so far there does not seem to be strong selective pressure for resistant parasites across the continent (Uwimana *et al*., 2020; Balikagala *et al*., 2021; Rosenthal, 2021; Rosenthal, Asua and Conrad, 2024). This suggests these mutants are being outcompeted in many areas, possibly due to a combination of higher fitness costs and different genetic backgrounds, alongside differences in competition between parasites (Cohen *et al*., 2012; Wasakul *et al*., 2023). For example, elevated transmission rates, combined with lower levels of drug pressure may increase the disadvantage generated by fitness costs (Abebaw *et al*., 2022; Agaba *et al*., 2022; Bylicka-Szczepanowska and Korzeniewski, 2022). This is reflected in African countries where ART-R is relatively more common, such as Eritrea. In some cases, these are areas which have made substantial reductions in total malaria burden and reduced transmission, both of which may mean the fitness costs of resistance mutations are less important, giving resistant genotypes a larger advantage (Fola *et al*., 2023). This could mean fitness costs play a role in the selective pressure for resistant variants, and although local-context specific, this could slow the emergence and spread of ART-R in some areas of Africa.

However, it is important to note that the role of fitness costs and genetic background in determining population level parasite dynamics is mostly speculative. To date, many of these effects have only been established *in vitro*, and in *in vivo* settings these costs likely interact with many other important factors which affect the relative competition between parasites, including drug use, vector dynamics, and other features of the parasite, such as antigenicity (Cohen *et al*., 2012; Myers-Hansen *et al*., 2020; He, Chaillet and Labbé, 2024). Moreover, even if these factors do play a role, it is clearly still possible for ART-R to emerge independently on divergent genetic backgrounds, and can be associated with varying degrees of selective pressure depending on local conditions, as seen in Southeast Asia (Takala-Harrison *et al*., 2015). Future studies should work toward combining genomic surveillance of parasites with *in vitro* and population level estimates of fitness costs, as this may allow for improved modelling of their role in determining epidemiological dynamics. This should be combined with whole-genome sequencing, as this would allow improved understanding of the role of genetic background on ART-R dynamics. If the importance of these factors could be established, it could be crucial for genomic surveillance, as it could affect the relative priority given to tracking cases of specific mutations/genetic backgrounds and their spread between areas (Mayor *et al*., 2023).

### 3.3 Rest of the World

We identified many *kelch13* mutations circulating across the rest of Africa, in particular West and Central Africa (Figure 7). For example, WHO validated/candidate mutations were detected at very low frequencies in Ghana, Mali, Nigeria, Equatorial Guinea, and the DRC (Figure 7). However, the majority of mutations in West and Central Africa did not have WHO validated/candidate status, and therefore have unclear phenotypic effects. For instance, non-WHO validated/candidate *kelch13* mutations were detected in Benin and Burkina Faso in West Africa, and in the Central African Republic, Chad, the DRC, and Equatorial Guinea In Central Africa (Figure 7). While the phenotypic effects of these mutations is unclear, some may indeed affect phenotypic resistance. We observed a mutation, R539I, at the same *kelch13* position as the WHO-validated R539T, making up a moderate proportion of the mutations in Togo, Ghana, Mali, and the DRC (Figure 7). Such a mutation should be surveilled closely and assessed in phenotypic assays of ART-R due to its position at a known resistance locus in *kelch13*. It is possible that different evolutionary pressures in West / Central Africa favour alternative substitutions at position 539 to Eastern Southeast Asia, where R539T is found.

**Figure 7.**
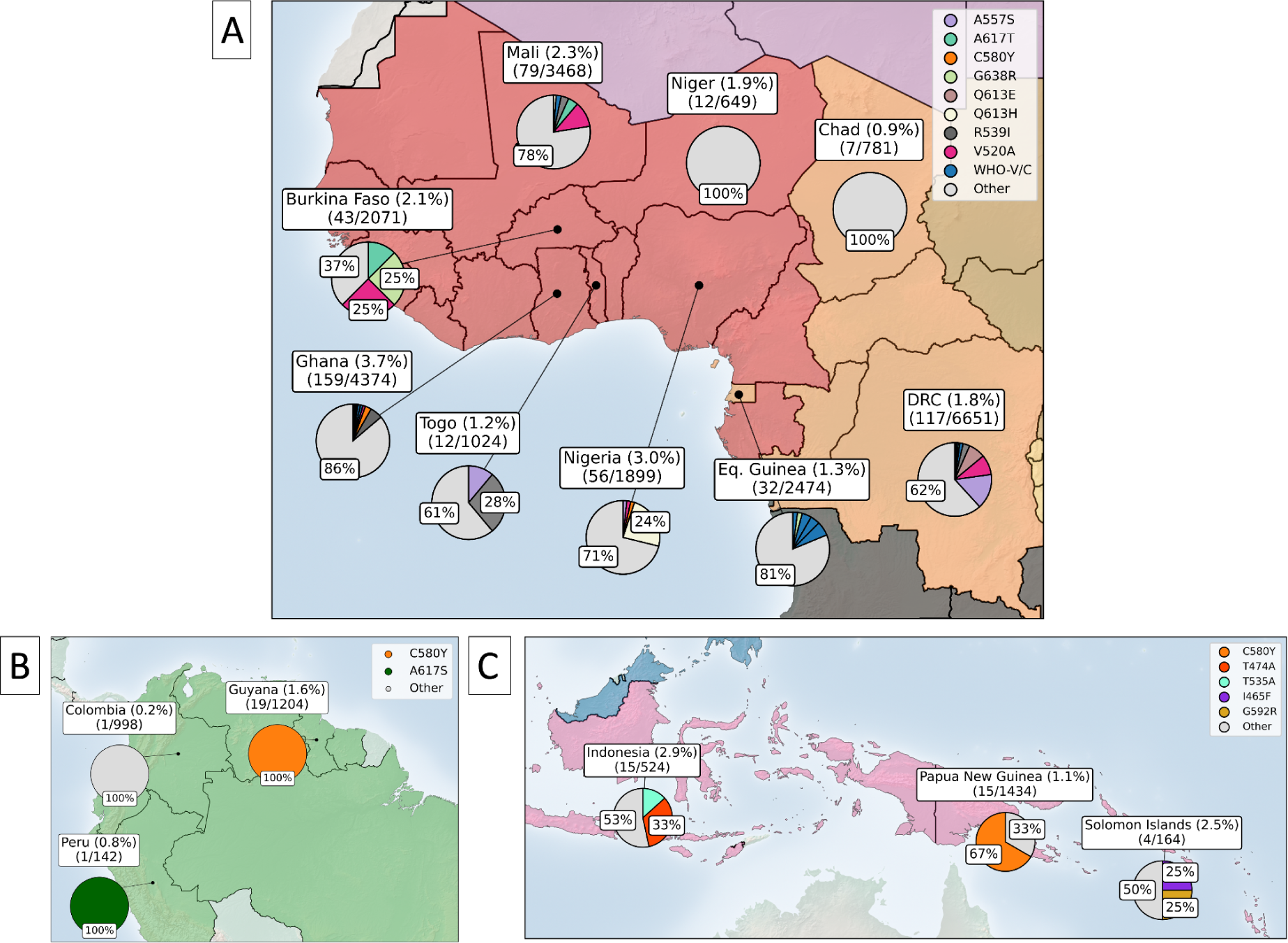
Distribution of *kelch13* propeller domain markers across West and Central Africa (A), South America (B) and Oceania (C). The percentage of samples with any *kelch13* mutation are included in the country labels, above the number of samples with any *kelch13* mutation, and the total number of samples collected for each country respectively. Among only the samples with any observed *kelch13* propeller domain mutation, pie charts show the proportions with markers of interest, where proportions above 25% are labelled accordingly. ‘Other’ denotes several low frequency mutations in the propeller domain which are not WHO validated/candidate markers, pooled into a single category.

Outside of Africa, one WHO validated/candidate mutation, C580Y, was detected in 19 samples from Guyana, South America (Figure 7). Aside from this, only two more mutations were detected In South America, each in one sample. In Peru, A617S was detected in 2016 and A504T was found in Colombia in 2018 (Figure 7). In Oceania, *kelch13* mutations were present in Indonesia, Papua New Guinea, the Solomon Islands, and Vanuatu. Evidence of ART-R was present in Papua New Guinea, where 67% of 15 samples with a mutation had C580Y. Another mutation circulating at relatively high frequencies within Oceania was T474A in Indonesia. In total, 25 non-WHO validated/candidate markers have been detected in Oceania between 1998 - 2020. WHO validated/candidate markers A675V, F446I, R539T, and R561H were also present in India (Figure 5). These markers are all shared with locations across Southeast Asia and A675V and R561H are also found in East/Northeast Africa (Figures 5 and 6) and highlight the potential for ART-R in India. ART-R does not appear to be taking hold in Far-East South Asia, where five non-validated/candidate mutations were detected in only five samples (0.002% of population). Similarly, in Western Asian countries of Afghanistan, Iran, and Pakistan very low proportions of mutant samples were found and all mutations were non-WHO validated/candidate (< 0.2% of population samples).

## 4. Predicting the next stage of ART-R in Africa

Given the emergence of ART-R in Africa, one obvious question is: what are the most likely scenarios for its spread? With the lessons learned from ART-R in Southeast Asia, it is useful to consider potential future scenarios on how ART-R might emerge and spread in Africa within the near future. Here, we articulate three possible scenarios regarding the frequency of *kelch13* mutations in Africa: a spread of similar magnitude to that of Southeast Asia, and two more optimistic scenarios - with or without additional interventions. It is possible that we may see different combinations of these situations play out across regions. We then discuss broader trends we might observe going forwards, including the human cost of ART-R in Africa, the effect of mitigation efforts on ART-R, regional variation in ART-R, and its dynamics over the next few years. In doing so, we hope this will inform long-term strategic thinking on mitigation strategies, and in properly allocating resources for future surveillance. It is important to note that any scenarios we outline are largely speculative, and need to be continually updated with new data as the situation progresses.

### 4.1.1 Scenario 1: Continued or increased spread of ART-R across Africa, similar to that seen in Southeast Asia

The emergence and spread of ART-R throughout Southeast Asia provides an example of how the situation might develop in Africa. In Southeast Asia, ART-R emerged in several independent locations, before a single drug resistant lineage spread across the region (Imwong, *et al*., 2017; Hamilton *et al*., 2019). The C580Y mutation then reached near-fixation (∼86%) in parts of Cambodia within a period of around 12-16 years (Figure 5) (Noedl *et al*., 2008; Dondorp *et al*., 2009; Kagoro *et al*., 2022). This offers one plausible scenario as to how ART-R might progress in Africa. It should be noted a worse possible scenario would be the spread of ‘full’ (as opposed to ‘partial’) artemisinin resistance, but since this has not been reported in the field, a more likely scenario is the continued spread of ART-R. As of 2024, this ‘partial’ resistance has now emerged independently in at least Uganda, Tanzania, Rwanda, Sudan and Eritrea (Uwimana *et al*., 2020; WHO, 2022; Conrad *et al*., 2023). Importantly, there is evidence of an increase in the proportion of samples with mutant *kelch13* markers in several of these countries since 2014, with similar increases in frequency to those observed in Southeast Asia around 10-15 years ago (Figure 8). If these areas were to follow a similar trajectory to areas of Cambodia for example, we could see frequencies of >54% delayed clearance under ACT treatment by around 2030 (Slater *et al*., 2016). Concerningly, this is the scenario highlighted by estimates of selection coefficients in Uganda, which suggest similar selective pressures to those observed in Southeast Asia between 2003-2018. These estimates imply the frequency of mutant *kelch13* parasites could reach as high as 95% between 2028-2033 (Meier-Scherling *et al*., 2024). Modelling also suggests that under the continued use of ACT therapies, *kelch13* mutations could become established in some areas (up to 57-88% frequency depending on model assumptions), though this will depend on their initial frequency and could potentially be reduced using TACT strategies (see Box 2) (Nguyen, Tran, *et al*., 2023). These increases in ART-R would in turn increase selective pressure for resistance to partner drugs, raising the likelihood of overall treatment failure. With more widespread ART-R, the pressure on partner drugs to clear infections is increased. Indeed, there is evidence of treatment failure potentially occurring in some areas of Angola and the Democratic Republic of the Congo (Dimbu *et al*., 2021; Moriarty *et al*., 2021; WHO, 2022a). Were such failures to become widespread, the human and economic consequences for the region would be disastrous (see Section 4.2.1). Unfortunately, the observed increases in ART-R across East and Northeast Africa in recent years suggest this scenario is fast becoming a reality.

**Figure 8.**
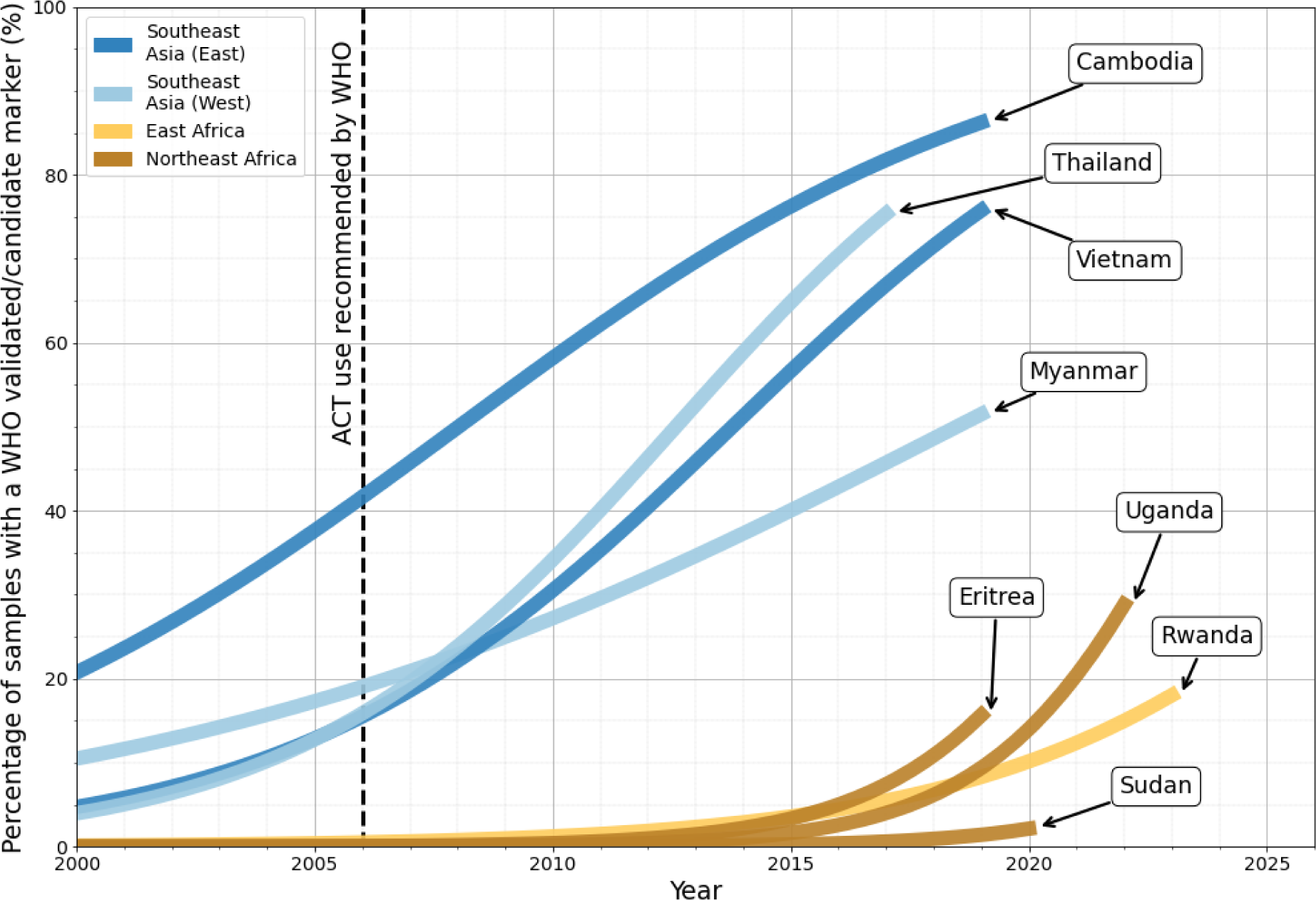
Proportion of samples with a WHO validated/candidate *kelch13* mutations over time (Table 2). Each country is shown using a logistic regression curve fitted to the observed data (see Figure S3 for model fits). Timepoints with no line indicate years with no available data. The black dashed line highlights 2006 - the year in which the WHO recommended ACT therapy as a first-line treatment measure in Africa (WHO, 2006). Notably, the rapid increase in prevalence of WHO *kelch13* markers in East and Northeast African countries is similar to that observed in Southeast Asian countries between 2007 and 2016. However, it is also important to note these trends have so far only been observed in a few countries in the East/North East of Africa and are based on smaller sample sizes (Assefa, Fola and Tasew, 2024).

### 4.1.2 Scenario 2: ART-R is limited in Africa by additional control interventions

It is clear that a massive and coordinated effort is urgently required to prevent the worst effects of ART-R from being realised across the African continent (Dhorda *et al*., 2024). This will require several mitigation strategies to be implemented to reduce the emergence and spread of ART-R. These strategies include the wide-scale implementation of Triple Artemisinin Combination Therapies (TACT) (see Box 2), or possibly Multiple Frontline Treatment (MFT) strategies, which modelling suggests could reduce treatment failure by 49-92% over a period of 5 years (Nguyen *et al*., 2015; Zupko *et al*., 2023; Li *et al*., 2024). If these mitigation strategies were implemented quickly and effectively, it may be possible to limit mutant *kelch13* to low frequencies in most areas, though regions where ART-R has already increased in frequency, such as Uganda, may see smaller reductions in the spread of resistant parasites. This could provide much needed time to develop additional antimalarial treatments (Phyo and von Seidlein, 2017; Siqueira-Neto *et al*., 2023). These strategies could be combined with the recently approved RTS,S and R21/Matrix-M vaccines, which in addition to improved vector control and case management, could reduce transmission chains in regions of high ART-R (WHO, 2022a; Datoo et al., 2024; Manzoni et al., 2024). However, while these strategies will likely reduce overall malaria burden, it should be noted they may actually promote resistance in the short term (see Section 4.2.2) (Wasakul *et al*., 2023).

It is also important to note the success of these strategies may also depend on the speed with which they are implemented. For example, introducing TACTs after ACT resistant parasites have reached even low frequency would likely have less of a preventative effect, with Nguyen *et al*., (2023) estimating this frequency could be as low as ∼1%. Moreover, the size and scale of parasite populations in Africa means any interventions will be complex and time-consuming, giving ART-R mutations a chance to rise to high frequency before mitigations are in place (WHO, 2022a). It is therefore a race against time to implement these strategies as quickly and effectively as possible, both to minimise the spread of ART-R and reduce overall malaria burden (Dhorda *et al*., 2024). Moreover, given progress in reducing overall malaria incidence in Africa has mostly plateaued since 2015, any single strategy is unlikely to reduce overall incidence to a level where widespread ART-R would not have dire consequences (Dhorda *et al*., 2024). It will therefore be crucial to combine mitigation strategies, in combination with careful genomic surveillance of parasite populations, as it may be possible to minimise the short term spread of ART-R by tailoring interventions to local conditions.

### 4.1.3 Scenario 3: Lower frequency and slower spread of ART-R in Africa than Southeast Asia, combined with additional interventions

While Scenario 1 and 2 are more likely scenarios, there are some reasons to think ART-R in Africa may not reach as high frequencies as Southeast Asia, or that if it were to, it may at least occur more slowly (Rasmussen, Alonso and Ringwald, 2022). For example, some areas of Southeast Asia, such as Myanmar, showed a slightly lower overall frequency and slower spread of ART-R than Cambodia and Thailand respectively, with several *kelch13* mutations other than C580Y reaching high frequency (Figure 5) (Tun *et al*., 2015; Kagoro *et al*., 2022). One possibility is that differential fitness costs or genetic backgrounds have slowed the spread of ART-R in these regions. Another possibility is that ecological factors, such as relatively higher rates of host immunity, asymptomatic infection, and counterfeit drugs, mean there is reduced overall drug use and selection for ART-R in these areas (Tun *et al*., 2015; Ataíde *et al*., 2017). For example, in East Africa, the presence of counterfeit artemisinin-based drugs has been confirmed. In Uganda, one study found ∼37% of 57 stores to be selling counterfeit ACTs (Björkman Nyqvist, Svensson and Yanagizawa-Drott, 2022) while anecdotal evidence of fake artesunate exists from Tanzania and Cameroon (Newton *et al*., 2006). If counterfeit ACTs contain insufficient doses of artemisinin, this is likely to increase selections for ART-R. However, if counterfeit ACTs containing no artemisinin are widespread in Africa, this may also lessen selection for ART-R, though of course at the cost of effectively treating patients with malaria. The role of genetic backgrounds, fitness costs, and ecological factors in slowing ART-R in Africa remains speculative (Conrad and Rosenthal, 2019; Rosenthal, Asua and Conrad, 2024). If these factors do play a role, their effects may be larger in African contexts compared with Southeast Asia, both because the effects of immunity and competition are more pronounced, and because the parasite population and geographical area in which resistance needs to spread are much larger, slowing its spread. For example, many areas of Africa are known to have higher rates of asymptomatic infection, population immunity, and strong competition between parasites, meaning the prevalence of ART-R may be reduced further relative to Southeast Asia (Cohen *et al*., 2012; Abebaw *et al*., 2022; Agaba *et al*., 2022; Bylicka-Szczepanowska and Korzeniewski, 2022).

While it is difficult to predict the precise effect fitness costs, alongside differences in transmission or genetic background might have, if they do play a role, one possible scenario could be ART-R frequencies in Africa similar to regions of Southeast Asia which have lower prevalence, such as Western and South Myanmar (Imwong, Dhorda, Myo Tun, *et al*., 2020; Kagoro *et al*., 2022). In such a scenario we may regularly see moderate frequencies of variant *kelch13* markers, which could reach much higher frequencies in some regions, depending on local conditions (WHO, 2022). This would result in ART-R being relatively common, though this may not be the norm across the continent. Nevertheless, this would still put increased pressure on partner drugs, increasing selection for resistance and increasing the likelihood of treatment failure. These factors could also interact with mitigation strategies, which in some cases could negatively affect their rollout or overall effectiveness. It will be necessary to conduct careful genomic surveillance to estimate the effects of these trends on ART-R.

### 4.2 Broader predictions on ART-R in Africa

#### 4.2.1 Widespread ART-R in Africa would be associated with significant human and economic costs

Given limited alternatives to ACT treatment, any increase in ART-R which might lead to increased partner drug failure would be catastrophic for human mortality in the region (Conrad and Rosenthal, 2019). While the scale of the problem has been widely acknowledged, few studies have quantified the human and economic impacts of widespread resistance (Lubell *et al*., 2014; Slater *et al*., 2016; WHO, 2022a). Modelling work suggests highly negative outcomes, depending on the frequency of artemisinin/partner-drug resistance (WHO, 2022a). For example, Lubell et al. (2014) found widespread ART-R (a conservative estimate of ∼30% ACT failure rate) could cause an additional 116,000 deaths annually, with medical costs of US$32 million. Moreover, decreased productivity could be over US$385 annually during the remaining lifetime of ACTs as the first-line treatment (Lubell *et al*., 2014). A second model considered several possible scenarios for ART-R and partner drug resistance in Africa, if resistance reached similar prevalence to various provinces in Southeast Asia (Slater *et al*., 2016). This model suggested if ART-R and partner drug resistance in Africa reached levels seen in Oddar Meanchey province in Cambodia (∼54%), this could result in around 16 million additional cases per year (∼7% overall increase) due to delayed parasite clearance and parasite recrudescence (Slater *et al*., 2016). Were this to take place, this could mean an additional 20,000 deaths and an economic impact of US$1.1 billion per year due to increased morbidity and mortality (Slater *et al*., 2016; Hailu *et al*., 2017; WHO, 2022). Unfortunately, these models offer conservative estimates and may even underestimate the true severity of the situation. The impact could be significantly worse if additional factors, such as diminished economic growth or lower educational outcomes were taken into account (Lubell *et al*., 2014; Slater *et al*., 2016; WHO, 2022a).

#### 4.2.2 The drive towards malaria elimination may promote resistance in the short term

Given the potential scale of the problem, national and international public health agencies have developed extensive plans to reduce total malaria burden (WHO, 2021, 2022a). For example, the World Health Organisation has stated their goal to eliminate malaria in 35 countries by 2030 (WHO, 2021, 2022a). While their success so far has been debated (Rosenthal, John and Rabinovich, 2019; WHO, 2022), as the total number of cases decreases, it is likely the proportion of drug resistant parasites will increase (White, 2004; Scott *et al*., 2018). This is for two reasons; firstly, since ACTs are a main strategy in treating infection, and ACT use selects for resistant parasites, resistant parasites are more likely to increase in the population (White, 2004). Secondly, as the overall parasite population and rate of infection is reduced, this will decrease immunity within the human population (Ataide *et al*., 2017; Agaba *et al*., 2022). As this occurs, the rate of asymptomatic infections (currently the most common kind of infection in many parts of Africa) may decrease, meaning more need for drug treatment and a further increase in selection for resistance (Bylicka-Szczepanowska and Korzeniewski, 2022).

This process is already being observed in some countries in Africa, such as Eritrea (Fola *et al*., 2023, Mihreteab Selam *et al*., 2023). Over the past two decades, concerted public health efforts, such as vector control, ACT treatment, and case management, have reduced infections in this region. However, some evidence suggests the success of these strategies means the proportion of ART-R parasites has increased, despite a reduction in overall burden (Fola *et al*., 2023, Mihreteab Selam *et al*., 2023). This trend has also been observed in several parts of Southeast Asia, such as Myanmar (Imwong *et al*., 2017b, 2020; Manzoni *et al*., 2024). This underscores the importance of careful genomic surveillance, as it can be used to identify patterns of selection on resistant parasites (Meier-Scherling *et al*., 2024). This would in turn inform mitigation strategies and help ensure that as overall parasite numbers are reduced, they are not replaced by resistant parasites.

#### 4.2.3 The prevalence of ART-R is likely to vary substantially between regions

A consequence of variation in reducing malaria burden is that the frequency of ART-R is also likely to be highly variable across the African continent, particularly over the next 1-5 years. This is not only because different African countries currently have very different malaria burdens, but are likely to have different rates of success in eliminating the disease going forwards (WHO, 2022). This variable distribution is also observed in Southeast Asia, where some regions, such as districts of Cambodia, show near fixation of mutant *kelch13* haplotypes while others, such as West and South Myanmar, have less than 20% prevalence (Kagoro *et al*., 2022). Unfortunately, the consequence is that some regions could become hubs for the spread of ART-R into other areas (Ashley *et al*., 2014; Tun *et al*., 2017). A similar effect was also observed in other regions of Southeast Asia, where several waves of selection allowed resistant lineages to spread across the region (Ashley *et al*., 2014; WHO, 2016; Tun *et al*., 2017). There are concerns these patterns could re-occur among African countries, where resistance-associated *kelch13* markers have so far only been observed in specific regions, such as Uganda, Rwanda and Eritrea (Uwimana *et al*., 2020; Agaba *et al*., 2022; Conrad *et al*., 2023). So far, it is not possible to determine whether a clonal spread of resistant genotypes is spreading in all of these countries due to the limited number of samples, though evidence suggests that so far these instances have emerged independently (Uwimana *et al*., 2020; Agaba *et al*., 2022; MalariaGEN *et al*., 2023; Conrad *et al*., 2023). This again highlights the importance of genomic surveillance and rapid data sharing, as when migration routes do arise, these can only be identified by monitoring and sequencing parasite populations, and used to inform mitigation strategies (Tun *et al*., 2017).

#### 4.2.4 The dynamics we observe in the next few years will determine the prevalence of ART-R in 10 years’ time

The patterns of resistance we see over the next 1-5 years will be crucial in determining the patterns we observe over the next decade. This is for four main reasons; firstly, the areas in which resistance first emerges gives indications of the areas in which resistance is currently under strong selection, and therefore likely to reach high frequencies. This effect was observed in Southeast Asia, where the regions in which resistant parasites initially emerged gave a strong indication as to where they would later reach high frequency, such as Cambodia and parts of Thailand (Imwong, *et al*., 2017; Kagoro *et al*., 2022). Secondly, resistance frequencies may reach critical thresholds in some areas within the next 1-5 years, beyond which mitigation strategies will be more difficult. For example, modelling studies have suggested that once the frequency of ACT resistant mutants reaches a critical threshold within a population, both MFTs and TACTs strategies are less likely to be effective (though this threshold can be as low as 1%) (Nguyen, Gao, *et al*., 2023; Zupko *et al*., 2023). Thirdly, these years will give a strong indication whether fitness costs and genetic background have a slowing effect on the spread of resistance in Africa, and this could assist in making predictive models for further spread. Lastly, we may see the initial effects of RTS,S and R21/Matrix-M vaccines during the next 1-5 years, and we can observe how this might affect selective pressures on the parasite (Datoo *et al*., 2024). Unfortunately, this period is also the time in which it is most difficult to alter mitigation strategies, simply because their planning and implementation takes time and resources from public health organisations. The next 1-5 years will therefore be where genomic surveillance and rapid data sharing is most crucial, as this could provide a warning system to identify at-risk areas and provide indications on how to best mitigate the spread of resistance. Incorporating whole genome sequencing into routine sampling will be particularly important, as this would enable discovery of new markers of resistance and detection of signatures of genomic adaptation to antimalarials, such as clonal population expansion. Whole genomes would also offer improved inferences on parasite dynamics, such as migration, population structure, and selection pressure, all of which will be crucial in tracking the spread of resistance.

## 5. The role of genomic surveillance going forward

Throughout this review, we have underscored the vital role genomic surveillance has played in understanding the global emergence and spread of ART-R in *P. falciparum*. In Southeast Asia, genomic surveillance helped to identify instances of migration of resistant parasites between countries and to quantify the frequencies of resistance in different areas (see Section 3). Now, in East and Northeast Africa, genomic surveillance continues to demonstrate its usefulness, by identifying the independent emergence and spread of *kelch13* mutations in Sudan, Uganda, Eritrea, and Rwanda (Mohamed *et al*., 2020; Uwimana *et al*., 2020; Owoloye *et al*., 2021; Conrad *et al*., 2023; Jalei *et al*., 2023; Mihreteab Selam *et al*., 2023). These discoveries were only possible due to the enhanced accuracy of genomic surveillance in detecting lower frequency genetic changes, as several TES from the region suggested effectiveness had remained mostly unchanged (>95% efficacy) (Assefa, Fola and Tasew, 2024). Although, TES primarily aid in understanding treatment failure due to partner drug resistance, so are not expected to inform on ART-R in the same way as genomic surveillance is able to do. Moving forward, it is essential to use the information gathered from both genomic surveillance and TES, along with lessons learned from ART-R in South-East Asia, to develop and implement mitigation strategies, such as properly allocating healthcare resources (WHO, 2015b, 2021; Assefa, Fola and Tasew, 2024).

### 5.1 The need for a more systematic approach to genomic surveillance

Genomic surveillance of ART-R in Africa is at a crucial inflection point. As of 2024, limitations on resources, time, and training, alongside the scale of parasite populations in Africa, means surveillance efforts remain localised, often with limited numbers of samples from key areas (Nsanzabana, 2021; WHO, 2022). This is a problem because surveillance is most valuable as a warning system when it is detailed enough to identify low frequency variants (e.g. less than 1%), and these can easily be missed if not enough samples have been analysed (Asua *et al*., 2019; Uwimana *et al*., 2020; Boni, 2022; Mayor *et al*., 2023). This highlights a fundamental challenge for the surveillance of ART-R, in that if resistance is circulating at lower frequencies, it is simply harder to detect using any method, and quantifying changes in those frequencies can be difficult without detailed sampling over time within the same areas (Mayor *et al*., 2023). While identifying resistance mutations at frequencies of around 5% or more is still very valuable, detecting mutations at lower frequencies gives more time in which to delay the establishment of resistant parasites in an area or their spread to neighbouring regions. These issues mean increases in resistance mutation frequencies may only be identified when they have effects on clinical efficacy (Nguyen, Gao, *et al*., 2023; Zupko *et al*., 2023).

A significant increase in genomic surveillance is needed, alongside a more systematic approach to sample collection. Surveillance could be improved by using networks of health facilities which routinely monitor ART-R frequencies within regions and conduct longitudinal sampling of areas over time (WHO, 2009; Nsanzabana *et al*., 2018; Guillot *et al*., 2022; Mayor *et al*., 2023). Adapting sampling for more novel surveillance approaches, including genomic surveillance, could help to maximise the available information on resistant allele frequencies and improve the detection of low-frequency variants (Mayor *et al*., 2023). An example of such an approach may involve performing *a priori* power analyses to guide sampling efforts, based on predefined estimates of mutation prevalence and desired margins of error (Mayor *et al*., 2023). More generally, the complementary use of whole genome sequencing and amplicon sequencing will be crucial to best tackle ART-R. While amplicon sequencing is likely to continue to be the most accessible genomic sequencing method in the near future, we must strive to use whole genome information to inform and improve amplicon sequencing. For example, whole genome sequencing can shed light on the “unknown unknowns” of ART-R, such as *de novo* drug resistance mutations, including those outside of the *kelch13* gene, which can then be incorporated into amplicon sequencing panels (Cerqueira *et al*., 2017). Aside from improving amplicon sequencing, whole genome data can also provide insights to partner drug resistance genotypes and parasite population dynamics (Parobek *et al*., 2017; Amato *et al*., 2018). Such information could inform models which characterise selective pressures in response to interventions like drug usage (Nguyen *et al*., 2023). Whole genome data can also highlight important changes in genomic architecture which can be indicative of evolution in response to control measures (Miotto *et al*., 2015). For example, identifying copy number variations in drug resistance genes, or deletions and breakpoints which lead to rapid diagnostic test failures (Rocamora and Winzeler, 2020). Moreover, increasing overall whole genome sequencing data and making it publicly available as soon as possible after collection would ensure maximum value from surveillance efforts.

Lastly, a major focus should be on building sequencing and data processing capabilities in malaria endemic countries, particularly in the African continent. Currently, the substantive cost of reagents, lack of sequencing technologies and issues with supply chain logistics can hinder efforts for in-country data generation (Neafsey, Taylor and MacInnis, 2021; Hamilton *et al*., 2023). Additionally, lack of training in specialist fields such as bioinformatics, genomics, and data science limits the ability to turn sequencing data into interpretable findings (Ishengoma *et al*., 2019). Challenges exist around obtaining sustained funding for long and short-term data storage, while the difficulties of data compartmentalization, standardisation, and administrative burdens will also need to be addressed to best leverage ongoing genomic surveillance efforts. Moving forwards, it will be critical to expand surveillance by enabling the use of sequencing more widely in Africa (Ishengoma *et al*., 2019). These improvements could include nanopore long-read technology, which has been demonstrated to be well-suited to surveillance needs through its increased portability, lower up-front costs, and real-time data output (Hamilton *et al*., 2023). Integrating genomic surveillance into public health requires stakeholder buy-in at all levels, thereby enabling policies and regulatory mechanisms. Several combined strategies could vastly enhance the region’s ability to respond to future outbreaks, both for *Plasmodium* and other pathogens. These strategies include systemic investment in local surveillance infrastructure and workflows, changes in sampling strategy (towards a framework informed by genomic surveillance), and a focus on building a critical mass of expertise in areas where gaps have been identified.

### 5.2 Improved models to analyse large spatiotemporal datasets

In addition to enhanced genomic surveillance, developing new models to analyse the increasingly large spatiotemporal datasets generated is crucial. Improved epidemiological models could characterise outbreaks as they unfold or even forecast how they may progress. For example, models which can estimate identity-by-descent among large numbers of parasites would enable inferences on population scale changes in genetic diversity (Miotto *et al*., 2013; Henden *et al*., 2018; Amambua-Ngwa *et al*., 2019). This capability would enhance quantification of selective pressures for ART-R, identify migration pathways between regions, and help categorise population diversity (Amato *et al*., 2018; Amambua-Ngwa *et al*., 2019; Meier-Scherling *et al*., 2024). Improved models could also study changes in the frequency of drug resistant lineages and facilitate the interpretation of these changes in relation to other factors, such as drug use, vector populations, or climate change (Ryan, Lippi and Zermoglio, 2020; Sherrard-Smith *et al*., 2022; Datoo *et al*., 2024; Redding, Gibb and Jones, 2024). Moreover, developing methods to identify novel resistance markers is crucial, for both *kelch13* and partner drug resistance genes, especially since resistance-conferring genetic mechanisms beyond *kelch13* have been identified in Africa, and potentially interact with the genetic background of African parasites (Miotto *et al*., 2015; Demas *et al*., 2018; Tumwebaze *et al*., 2022). This will be particularly important during the rollout of MFT or TACT strategies, as any triply resistant mutants which arise can be more easily identified using genomic surveillance, and this information could inform optimal timing to rotate drugs (Zupko *et al*., 2023; Kokori *et al*., 2024, Boni, 2022; Nguyen *et al*., 2023).

### 5.3 Improved data sharing, public access to data and training opportunities

Ensuring that surveillance data and analysis methods are publicly available is crucial for the malaria community, as these resources are essential in building skills and capacity for bioinformatic analysis. Public access to large surveillance datasets has repeatedly empowered novel analyses on the basic biology of *Plasmodium*, possible only because of the open nature of these datasets and the willingness of the community to share data (MalariaGEN *et al*., 2023). For example, recent analysis of the Pf7 open dataset of whole genome sequencing samples led to the identification of a novel ‘cryptotype’ of *P. falciparum*, present in 13 countries across Africa (Miotto *et al*., 2024). Additionally, making the outputs of genomic surveillance data available through centralised platforms like MalariaGEN, WWARN, and WHO’s malaria threat maps has provided the malaria community with invaluable resources to study the distribution, migration and spread of ART-R (Sibley, Barnes and Plowe, 2007; MalariaGEN *et al*., 2023). Similarly, web applications like Pf-HaploAtlas have been developed to leverage these resources, facilitating analysis of genomic surveillance data without the need for specialised technical knowledge (Lee *et al*., 2024). It is imperative to continue this work by ensuring these tools and resources are accessible and user-friendly, particularly for those with limited training, and to support the malaria community in making their data publicly available as soon as possible after collection. This approach will enhance contributions and coordination within the community, and help in informing strategies to mitigate the spread of ART-R.

## 6. Conclusions and outlook

Given the emergence of ART-R associated with *kelch13* markers in Africa, it is likely only a matter of time before these begin to spread widely across the continent (Dhorda *et al*., 2024; Rosenthal, Asua and Conrad, 2024). This review has highlighted the crucial role of genomic surveillance in alerting public health agencies to the spread of ART-R and in planning the use of limited resources to contain it. Recent years have seen limitations on funding, political instability, and other public health priorities, such as the COVID-19 pandemic, divert already limited resources away from genomic surveillance of *P. falciparum*, jeopardising the surveillance of ART-R in Africa (Lyimo *et al*., 2022; Mensah, Akyea-Bobi and Ghansah, 2022). Despite these challenges, it will be crucial to continually develop and champion the use of genomic surveillance as a vital part of the surveillance toolkit for ART-R, which also contains methods such as treatment efficacy studies and *in vitro* phenotypic studies. Without coordinating these surveillance tools, increases in ART-R may only be identified when they have already spread widely, negatively affecting patient outcomes and compromising our ability to implement mitigation strategies (Nguyen, Gao, *et al*., 2023; Zupko *et al*., 2023). This means continuing surveillance is vital, not only for containing the spread of ART-R, but also for minimising the human costs it will cause.

## BOXES

### Box 1 Fitness costs associated with ART-R

Fitness costs could significantly impact the spread of ART-R. In the presence of the drug, resistant parasites have a selective advantage, making them more likely to transmit between individuals and spread within a population (White, 2004). However, if there are fitness costs to being resistant, then in the absence of the drug, resistant parasites would be outcompeted by susceptible genotypes, making them more likely to remain at low frequency (Rosenthal, 2013).

Numerous *in vitro* studies have demonstrated fitness costs associated with *kelch13* mutations, such as decreased parasite blood stage growth-rate (Straimer *et al*., 2017) and competitive growth disadvantage (Nair *et al*., 2018; Mathieu *et al*., 2020; Stokes *et al*., 2021). Delayed parasite clearance under artemisinin exposure is linked to decreased haemoglobin endocytosis, limiting the parasite’s ability to utilise important amino acids (Bunditvorapoom *et al*., 2018). *kelch13* mutant parasites also have lower heme levels during the trophozoite stage, which may cause growth defects (Heller and Roepe, 2018; Heller, Goggins and Roepe, 2018). *kelch13* mutations may also affect stability of the protein, leading to transcriptional stress responses and cell development issues (Mok *et al*., 2015; Yang *et al*., 2019; Behrens *et al*., 2023). Moreover, different *kelch13* mutations can confer differing levels of ART-R and fitness costs. For example, C580Y and R561H show similar increased survival under drug treatment, but differing fitness costs (Nair *et al*., 2018). The F446I mutation, common in Myanmar, provides a small survival advantage during drug treatment, but lower competitive growth *in vitro* than C580Y (Stokes *et al*., 2021). While other regions of the genome have also been associated with decreased clearance rate for artemisinin (Cerqueira *et al*., 2017; Mukherjee *et al*., 2017), these mechanisms are complex, and require further experimental validation of potential fitness costs (for a review, see Pandit *et al*., 2023).

The presence of these costs is unsurprising given the highly conserved nature of the *kelch13* propeller domains, yet may have important consequences for selection for ART-R (Ariey *et al*., 2014; Miotto *et al*., 2015). Fitness costs may be one reason why *kelch13* mutations are typically observed in isolation, rather than multiple mutations within a single parasite. This effect was seen in our dataset, where in a single study, only one double *kelch13* propeller mutation (M579T/N657H) occurred in more than 5 samples (Mishra *et al*., 2017). However, as neither M579T and N657H were found individually or together in other studies, this double mutation may have arisen from laboratory cross-contamination or mixed-strain infection.

### Box 2 What treatment strategies could slow the spread of treatment failure?

ART-R is a major concern because of its broader role in ACT treatment failure. In essence, ART-R increases selective pressure for partner drug resistance, which in turn increases the likelihood of ACT treatment being unsuccessful. Several strategies have been proposed to slow the spread of treatment failure, including ACT cycling, multiple first-line therapies (MFTs), and triple artemisinin combination therapies (TACTs). Each of these methods have been used at various times and locations, though all have varying costs, benefits, and degrees of evidence for their efficacy (Pluijm *et al*., 2021; Boni, 2022; Chen and Hsiang, 2022). It is important to note under each of these strategies, mutants which are resistant to both artemisinin derivatives and partner drugs may eventually arise (Figure 9). Having a strategy for if this occurs will be paramount, and will require careful genomic surveillance to identify cases where they emerge (Mayor *et al*., 2023).

**Figure 9.**
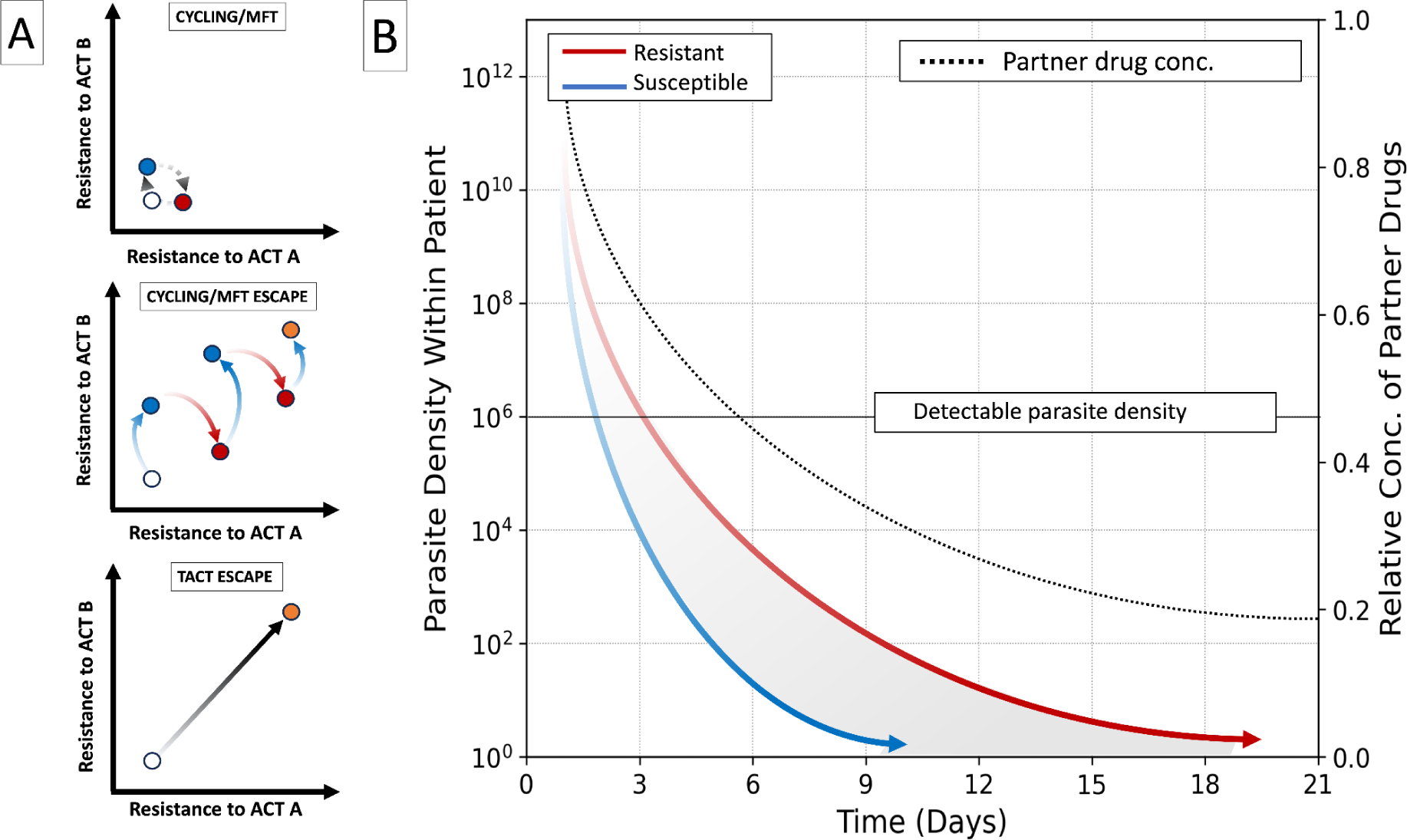
A) There are three possible evolutionary scenarios during cycling, MFT or TACT strategies. In the idealised cycling/MFT scenario (top panel), resistance will cycle between two states of elevated resistance to a given ACT combination therapy, but not both, at any given time (red/blue points). However, these strategies could promote dual resistant escape mutants through stepwise increases over time (middle panel). This may occur even if resistance to any individual ACT decreases in the short term (red and blue lines). In contrast, triple artemisinin-based combination therapies (bottom panel) may directly select for triple resistant escape mutants (black solid line). While the likelihood of this occurring may be lower than under cycling or MFT strategies, this could result in mutants which are resistant to multiple partner drugs, leaving limited options to change strategy (image adapted from Baym, Stone and Kishony, 2016). B) TACTs involve the use of two partner drugs throughout treatment. Here, a theoretical example of parasite density over time within a patient is shown for artemisinin resistant and susceptible parasites (red and blue lines respectively). In cases where parasites are resistant to artemisinin, their density remains higher after initial artemisinin exposure (∼ 3 days), resulting in more parasites for the partner drug to treat (shaded area), and therefore stronger selection for resistance to that drug. TACTs use two partner drugs (black dashed line), which in theory means selection for any individual drug is lower, and the chance of triply resistant parasites emerging within any given infection is less likely (Nguyen *et al*., 2023, image adapted from Hanboonkunupakarn and White, 2022).

#### ACT cycling

As of 2022, the WHO recommends a policy of ‘ACT cycling’, which involves using a single first-line ACT, before switching to a second once efficacy decreases below a certain threshold in the population (∼ 90%) (WHO, 2015a, 2022). Theoretically, this should reduce the frequency of resistance to the first drug, by providing time for resistance to decline in frequency while the second drug is in use (Figure 9) (Boni, Smith and Laxminarayan, 2008; Nguyen *et al*., 2015; Boni, 2022). This strategy initially proved effective in Cambodia, where artesunate-mefloquine was replaced with dihydroartemisinin–piperaquine, before reverting back after resistant parasites reduced in frequency (Ross *et al*., 2018; Rasmussen, Alonso and Ringwald, 2022). However, there is some evidence to suggest the strategy later drove the emergence of triply resistant mutants in the Northeast of the country (Rossi *et al*., 2017).

#### Multiple front-line therapies

More recently, several countries have adopted the use of MFTs, i.e. using multiple ACT therapies concurrently within a population (WHO, 2021). Importantly, modelling studies have found MFTs have similar efficacy to cycling strategies but are less likely to accelerate the emergence and spread of resistance (Boni, Smith and Laxminarayan, 2008; Smith *et al*., 2010; Nguyen *et al*., 2015). This work suggests MFTs are most effective when using partner drugs with different mechanisms of action, as this reduces the chances of cross-resistance to both drugs (Li *et al*., 2024). However, the WHO has stopped short of recommending MFTs more generally (WHO, 2013), citing contradictions in results from modelling work, and challenges in locations with both high drug use and resistance (Boni, 2022; Antao and Hastings, 2012). For example, the high frequency of the artemisinin resistant lineage in Cambodia means there is a lack of alternative treatments should MFTs fail (Ross *et al*., 2018; Rasmussen, Alonso and Ringwald, 2022).

#### Triple artemisinin-based combination therapies (TACTs)

Another alternative is TACTs, which combine an artemisinin derivative with two partner drugs. Theoretically, this method preserves treatment efficacy, because of the rarity of parasites acquiring resistance to both partner drugs over a course of treatment (Figure 9) (Pluijm *et al*., 2020). Recently, large-scale clinical trials in Southeast Asia have demonstrated TACTs are effective at national scales, even in situations where multidrug-resistant parasites are present (Pluijm *et al*., 2020; Peto *et al*., 2022). Modelling studies suggest that TACTs could significantly delay the emergence and spread of ART-R compared to other strategies (Nguyen *et al*., 2023; Zupko *et al*., 2023; Li *et al*., 2024). However, their effectiveness diminishes once resistant parasites reach a threshold frequency in the population, which can be as low as 1%. A further drawback of TACTs is that because three drugs in total are being administered, the frequency of negative side-effects on patient health is more common (Pluijm *et al*., 2020, 2021; Peto *et al*., 2022). While promising, for TACTs to be useful at population scales, several practical issues need to be ironed out, such as correct dosing and formulation of partner drugs for overall safety, alongside increased cost-effectiveness and regional availability (for a review, see Kokori *et al*., 2024). Moreover, their use requires public acceptance, as the reduced risk of emergent resistance is being balanced against an increased risk of side-effects for any individual taking the drugs (Pluijm *et al*., 2020, 2021).

## Supporting information

Supplementary data README

## Glossary

ART-R: Artemisinin partial resistance
ACT: Artemisinin Combination Therapy
RSA: Ring-stage Survival Assay
TACTs: Triple Artemisinin Combination Therapies
MFT: Multiple Front-line Therapies
WGS: Whole Genome Sequencing
WHO: World Health Organisation
Propeller mutations: non-synonymous mutations occurring in the BTB/POZ and propeller domains of the *kelch13* gene (codons 349 - 726)
AL: artemether-lumefantrine
AS: artesunate
AQ: amodiaquine
MQ: mefloquine
DHA: dihydroartemisinin
PYR: pyronaridine
CQ: chloroquine
PPQ: piperaquine
PQ: primaquine
SP: Sulfadoxine-pyrimethamine

