## Supplementary Information: Methods for "Understanding the Global Spread of Artemisinin Resistance: Insights from over 100K *Plasmodium falciparum* Samples"

### Supplementary Methods

#### Data Collection and Processing for Spatiotemporal Analysis of *kelch13* in 112,933 samples

We collected *kelch13* amino acid mutation data from three sources: the MalariaGEN Pf7 data release (MalariaGEN et al. 2023), the Worldwide Antimalarial Resistance Network (WWARN) Artemisinin Resistance Molecular Surveyor database (https://www.iddo.org/wwarn/tracking-resistance/artemisinin-molecular-surveyor), and our own literature search. Mutations were recorded alongside sample ID, continent, population, and country of origin, plus year of collection and publication source. In total, we collated data from 112,933 *Plasmodium falciparum* samples from 73 countries between the years of 1980 - 2023. The datasets, including per sample marker and per population marker count, are available as supplementary datasets S1 and S2.

Samples were allocated to one of thirteen populations. Ten populations were named according to the Pf7 dataset, where samples were grouped based on a geographical and genetic analysis (MalariaGEN et al. 2023). These are: South America, West Africa, Central Africa, Northeast Africa, East Africa, Eastern South Asia, Far-Eastern South Asia, Western Southeast Asia, Eastern Southeast Asia, and Oceania. To simplify analyses, three changes were made to Pf7 population definitions. In Pf7, Kenya, India, and Thailand are split across two populations. In this study, we assigned these countries to the population which held most samples. All samples from Thailand were allocated to Western Southeast Asia, all samples from Kenya were allocated to East Africa, and all samples from India were allocated to Eastern South Asia. Three extra populations were created for this analysis, for samples which did not fit into pre-existing populations geographically: Southern Africa (Angola, Mozambique, South Africa, Zambia, Zimbabwe), North Africa (Algeria, Libya), Western Asia (Afghanistan, Iran, Pakistan) (Table S2). Pf7 samples from Mozambique were re-allocated to Southern Africa. For a summary of countries and sample sizes within each population, see Table S2.

For all data sources we applied the following inclusion criteria to reported mutations:

1. Occurred within the *kelch13* BTB/POZ and propeller domains at codon positions 349-726. Samples with no *kelch13* mutations or with a mutation outside of these positions were categorised as having the 3D7 reference sequence. In some cases, samples with two *kelch13* mutations were identified. When one mutation was outside of the propeller domain, we counted only the propeller domain mutation for that sample. Where both mutations were detected within the propeller domain, we recorded both mutations for the sample, meaning it could be a mixed strain or polymutant.
2. Mutation induced a non-synonymous amino acid change. We included samples with only synonymous amino acid mutations as 3D7 reference.

We classified all *kelch13* mutations according to their status for association with ART-R from the latest WHO guidelines - either 'validated', 'candidate' or 'no status' https://www.who.int/news-room/questions-and-answers/item/artemisinin-resistance (Table S1) . Mutation A578S was classified as 'not associated' with ART-R (i.e. known to be artemisinin susceptible) (i.e. known to be artemisinin susceptible) (WHO, 2022).

Whole genome data from 16,203 QC-passed Pf7 samples were processed using custom Python scripts to generate haplotypes for *kelch13*. For each sample, we applied any nucleotide variants from the GT field of the sample's VCF file to the 3D7 reference sequence for *kelch13*. These sequences were translated into amino acids, and *kelch13* mutations were identified. We excluded any sample with a missing genotype call (n = 1,056). After removal of heterozygous samples (n = 2,292), 12,855 samples from Pf7 were added to the final dataset.

All *kelch13* data from WWARN's Artemisinin Molecular Surveyor were downloaded. We reviewed 63 studies where only a range of years, rather than a single year, was provided. If we were not able to gather single-year data for all samples from the original publication, we recorded the median year of the range for the study. In cases where the year range was two years, we selected the year with the largest number of samples. We examined the full texts of 34 WWARN publications that had a mismatched total number of mutations and total number of tested samples. Of these, 17 studies were included after correcting for recording errors and one was included after being flagged as containing multiple propeller mutations per sample. The remainder were excluded for containing heterozygous samples, or due to missing data for most samples, or because the publication was inaccessible. In total,In total, we included data from 291 WWARN publications covering 86,870 samples.

To ensure our database was as comprehensive as possible, we performed an additional literature search for relevant, recent papers published after 2020 to supplement the WWARN database. We searched the Web of Science and PubMed with the following string: "K13 OR kelch OR kelch13 OR pfkelch13) AND (half life OR parasite clearance OR resistance)", restricting results to articles only. We also searched "Kelch13 mutations falciparum artemisinin" in Google Scholar for articles published between 2023 - 2024. We excluded articles which genotyped *kelch13* after ACT treatment (i.e. not day 0 patients) or in cultured labs strains, articles which did not provide country-level information or sample size information, articles which only provided genotypes for a subset of samples tested, articles already included in the WWARN database, and articles not available in English or not accessible. We yielded 28 additional unique studies after reviewing full texts and applying both the article and mutation inclusion criteria. A further 13,208 samples were collected from this search.

#### Supplementary Tables and Figures

**Table S1.** Criteria for *kelch13* mutations to be classified as candidate or validated markers of artemisinin partial resistance by the WHO (WHO, 2022) and the current list of accepted mutations in each category.

| Classification | Criteria | List of current *kelch13* mutations |
|---|---|---|
| **Candidate or associated *kelch13* markers of artemisinin partial resistance** | 1. A statistically significant association ($p < 0.05$) between a *kelch13* mutation and clearance half-life >5 hours or day 3 parasitaemia via a chi-squared test or appropriate multivariable regression model on a sample of at least 20 clinical cases<br>**OR**<br>2. Survival of >1% using the $RSA^{0–3h}$ in at least five individual isolates with a given mutation or a statistically significant difference ($p < 0.05$) in the $RSA^{0–3h}$ assay between culture-adapted recombinant isogenic parasite lines, produced using transfection and gene editing techniques, which express a variant allele of *kelch13* as compared with the wild-type allele | P441L, G449A, C469F, A481V, R515K, P527H, N537I, N537D, G538V, V568G |
| **Validated *kelch13* markers of artemisinin partial resistance** | Both requirements 1 and 2 are met | F446I, N458Y, C469Y, M476I, Y493H, R539T, I543T, P553L, R561H, P574L, C580Y, R622I, A675V |
| RSA = ring-stage survival assay.This assay measures the survival of early ring-stage parasites after a six hour incubation with 700nM dihydroartemisinin. Abnormal RSA result = >1-2% survival compared to control 66 hours after incubation (Rosenthal et al. 2024). | | |

**Table S2.** The thirteen populations included in this analysis and the countries and sample sizes within each population. Ten populations match those outlined in MalariaGEN's Pf7 release and are guided by both geographical and genetic patterns while three extra populations were created for this analysis, for samples which did not fit into pre-existing populations geographically: Southern Africa, North Africa, and Western Asia. Proportion of population = the percentage of samples contributed by each country to its populations. Proportion of total = the percentage of samples contributed by each country to the total dataset. DRC = Democratic Republic of the Congo.

| Population | Country | Sample Count | Proportion of Population (%) | Proportion of Total (%) |
|---|---|---|---|---|
| South America | Brazil | 764 | 16.35 | 0.68 |
| | Colombia | 998 | 21.36 | 0.88 |
| | Dominican Republic | 8 | 0.17 | 0.01 |
| | Ecuador | 70 | 1.5 | 0.06 |
| | French Guiana | 977 | 20.91 | 0.87 |
| | Guyana | 1,204 | 25.77 | 1.07 |
| | Haiti | 162 | 3.47 | 0.14 |
| | Peru | 142 | 3.04 | 0.13 |
| | Suriname | 40 | 0.86 | 0.04 |
| | Venezuela | 307 | 6.57 | 0.27 |
| West Africa | Benin | 815 | 3.26 | 0.72 |
| | Burkina Faso | 2,071 | 8.3 | 1.83 |
| | Côte d'Ivoire | 2,296 | 9.2 | 2.03 |
| | Cameroon | 2,183 | 8.75 | 1.93 |
| | Cape Verde | 39 | 0.16 | 0.03 |
| | Gabon | 1,078 | 4.32 | 0.95 |
| | Gambia | 1,177 | 4.72 | 1.04 |
| | Ghana | 4,374 | 17.52 | 3.87 |
| | Guinea | 1,169 | 4.68 | 1.04 |
| | Guinea-Bissau | 19 | 0.08 | 0.02 |
| | Liberia | 437 | 1.75 | 0.39 |
| | Mali | 3,468 | 13.89 | 3.07 |
| | Mauritania | 75 | 0.3 | 0.07 |
| | Niger | 649 | 2.6 | 0.57 |
| | Nigeria | 1,899 | 7.61 | 1.68 |

| | | | | |
|---|---|---|---|---|
| | Sao Tome and Principe | 120 | 0.48 | 0.11 |
| | Senegal | 1,862 | 7.46 | 1.65 |
| | Sierra Leone | 207 | 0.83 | 0.18 |
| | Togo | 1,024 | 4.1 | 0.91 |
| Central Africa | Central African Republic | 201 | 1.95 | 0.18 |
| | Chad | 781 | 7.56 | 0.69 |
| | Congo | 222 | 2.15 | 0.2 |
| | DRC | 6,651 | 64.39 | 5.89 |
| | Equatorial Guinea | 2,474 | 23.95 | 2.19 |
| East Africa | Burundi | 1 | 0.01 | 0 |
| | Comoros | 627 | 5.15 | 0.56 |
| | Kenya | 4,265 | 35.06 | 3.78 |
| | Madagascar | 367 | 3.02 | 0.32 |
| | Malawi | 506 | 4.16 | 0.45 |
| | Rwanda | 2,154 | 17.71 | 1.91 |
| | Somalia | 325 | 2.67 | 0.29 |
| | Tanzania | 3,920 | 32.22 | 3.47 |
| North Africa | Algeria | 2 | 22.22 | 0 |
| | Egypt | 1 | 11.11 | 0 |
| | Libya | 6 | 66.67 | 0.01 |
| Northeast Africa | Djibouti | 9 | 0.08 | 0.01 |
| | Eritrea | 1,563 | 13.32 | 1.38 |
| | Ethiopia | 732 | 6.24 | 0.65 |
| | Saudi Arabia | 4 | 0.03 | 0 |
| | South Sudan | 323 | 2.75 | 0.29 |
| | Sudan | 1,123 | 9.57 | 0.99 |
| | Uganda | 7,752 | 66.04 | 6.86 |
| | Yemen | 232 | 1.98 | 0.21 |
| Southern Africa | Angola | 1,635 | 25.72 | 1.45 |
| | Mozambique | 1,103 | 17.35 | 0.98 |
| | South Africa | 2,926 | 46.03 | 2.59 |

| | | | | |
|---|---|---|---|---|
| | Zambia | 684 | 10.76 | 0.61 |
| | Zimbabwe | 9 | 0.14 | 0.01 |
| Western Asia | Afghanistan | 342 | 31.46 | 0.3 |
| | Iran | 76 | 6.99 | 0.07 |
| | Pakistan | 669 | 61.55 | 0.59 |
| Eastern South Asia | India | 4,261 | 100 | 3.77 |
| Far-Eastern South Asia | Bangladesh | 2,485 | 98.61 | 2.2 |
| | Nepal | 35 | 1.39 | 0.03 |
| Western Southeast Asia | China | 1,139 | 6.12 | 1.01 |
| | Myanmar | 11,868 | 63.79 | 10.51 |
| | Thailand | 5,599 | 30.09 | 4.96 |
| Eastern Southeast Asia | Cambodia | 6,213 | 44.26 | 5.5 |
| | Laos | 2,659 | 18.94 | 2.35 |
| | Malaysia | 342 | 2.44 | 0.3 |
| | Philippines | 213 | 1.52 | 0.19 |
| | Vietnam | 4,610 | 32.84 | 4.08 |
| Oceania | Indonesia | 524 | 23.93 | 0.46 |
| | Papua New Guinea | 1,434 | 65.48 | 1.27 |
| | Solomon Islands | 164 | 7.49 | 0.15 |
| | Vanuatu | 68 | 3.11 | 0.06 |

**Table S3**. Number of unique *kelch13* propeller mutations across populations. C/V = WHO candidate or validated *kelch13* marker associated with partial artemisinin resistance.

| Continent | Population | Total Samples | No. Mutations | No. C/V Mutations | % of samples with a mutation | % of samples with a C/V mutation |
|---|---|---|---|---|---|---|
| South America | SA | 4,672 | 3 | 1 | 0.45 | 0.41 |
| Africa | AF-W | 24,962 | 185 | 5 | 2.09 | 0.06 |
| | AF-C | 10,329 | 96 | 7 | 1.98 | 0.12 |
| | AF-N | 9 | 0 | 0 | 0 | 0 |
| | AF-NE | 11,738 | 81 | 7 | 10.39 | 8.08 |
| | AF-E | 12,165 | 108 | 11 | 3.81 | 1.45 |
| | AF-S | 6,357 | 42 | 4 | 0.98 | 0.09 |
| Asia | AS-W | 1,087 | 5 | 0 | 0.64 | 0 |
| | AS-S-E | 4,261 | 13 | 4 | 2.89 | 0.54 |
| | AS-S-FE | 2,520 | 6 | 0 | 0.52 | 0 |
| | AS-SE-W | 18,606 | 96 | 18 | 34.63 | 32.37 |
| | AS-SE-E | 14,037 | 41 | 13 | 51.59 | 50.82 |
| Oceania | OC | 2,190 | 24 | 1 | 1.74 | 0.46 |

**Table S4.** Proportion of total samples per year from each population.

| Year | South America | West Africa | Central Africa | North Africa | Northeast Africa | East Africa | Southern Africa | Western Asia | Eastern South Asia | Far-Eastern South Asia | Western Southeast Asia | Eastern Southeast Asia | Oceania | Total Samples |
|---|---|---|---|---|---|---|---|---|---|---|---|---|---|---|
| **1980** | 0 | 0 | 0 | 0 | 0 | 0 | 0 | 0 | 0 | 0 | 0 | 0 | 100 | 13 |
| **1984** | 0 | 100 | 0 | 0 | 0 | 0 | 0 | 0 | 0 | 0 | 0 | 0 | 0 | 69 |
| **1987** | 0 | 0 | 0 | 0 | 0 | 0 | 0 | 0 | 0 | 0 | 0 | 0 | 100 | 6 |
| **1990** | 0 | 100 | 0 | 0 | 0 | 0 | 0 | 0 | 0 | 0 | 0 | 0 | 0 | 8 |
| **1991** | 0 | 5.26 | 0 | 0 | 0 | 0 | 0 | 0 | 0 | 0 | 94.74 | 0 | 0 | 38 |
| **1993** | 0 | 0 | 0 | 0 | 0 | 0 | 0 | 0 | 0 | 0 | 0 | 100 | 0 | 5 |
| **1995** | 0 | 0 | 0 | 0 | 0 | 100 | 0 | 0 | 0 | 0 | 0 | 0 | 0 | 168 |
| **1996** | 0 | 0 | 0 | 0 | 0 | 51.19 | 0 | 0 | 0 | 0 | 0 | 0 | 48.81 | 84 |
| **1997** | 0 | 0 | 0 | 0 | 0 | 23.4 | 0 | 0 | 0 | 0 | 40.43 | 36.17 | 0 | 94 |
| **1998** | 0 | 0 | 0 | 0 | 0 | 63.93 | 0 | 0 | 0 | 0 | 0 | 0 | 36.07 | 305 |
| **1999** | 4.26 | 0 | 0 | 0 | 93.95 | 0.14 | 0 | 0 | 0 | 0 | 0 | 1.65 | 0 | 727 |
| **2000** | 0 | 0 | 0 | 0 | 0 | 78.31 | 0 | 0 | 0 | 0 | 0 | 21.69 | 0 | 83 |

| | | | | | | | | | | | | | | |
|---|---|---|---|---|---|---|---|---|---|---|---|---|---|---|
| **2001** | 0 | 14.47 | 64.15 | 0 | 0 | 0 | 0 | 0 | 0 | 0 | 21.38 | 0 | 0 | 159 |
| **2002** | 0 | 21.95 | 0 | 0 | 0 | 0 | 0 | 0 | 0 | 0 | 30.73 | 47.32 | 0 | 410 |
| **2003** | 6.04 | 37.74 | 0 | 0 | 0 | 7.55 | 9.43 | 0 | 0 | 0 | 25.28 | 2.64 | 11.32 | 530 |
| **2004** | 0 | 12.08 | 30.72 | 0 | 0 | 6.8 | 0 | 0 | 0 | 0 | 37.51 | 12.9 | 0 | 853 |
| **2005** | 0 | 5.71 | 9.19 | 0 | 8.08 | 24.56 | 0 | 0 | 0 | 0 | 37.08 | 15.37 | 0 | 631 |
| **2006** | 0 | 11.92 | 44.77 | 0 | 0 | 19.62 | 0 | 0 | 0 | 0 | 8.43 | 15.26 | 0 | 688 |
| **2007** | 0 | 22.62 | 13.23 | 0 | 3.17 | 16.05 | 0 | 0 | 0 | 1.56 | 41.44 | 1.92 | 0 | 2,237 |
| **2008** | 35.45 | 4.69 | 0 | 0 | 3.65 | 1.56 | 0 | 0 | 0 | 0.57 | 36.73 | 11.61 | 5.73 | 2,110 |
| **2009** | 0.74 | 23.2 | 0 | 0 | 0 | 9.43 | 0 | 0 | 6.77 | 3.46 | 29.6 | 25.77 | 1.03 | 1,358 |
| **2010** | 3.37 | 31.2 | 1.41 | 0 | 4.92 | 13.36 | 1.72 | 0 | 11.5 | 0 | 9.37 | 22.31 | 0.83 | 2,904 |
| **2011** | 0.51 | 12.3 | 0 | 0 | 4.45 | 11.28 | 0.15 | 0 | 3.34 | 7.37 | 14.31 | 45.77 | 0.51 | 5,276 |
| **2012** | 12.15 | 15.32 | 1.84 | 0 | 5.14 | 12.62 | 4.29 | 0 | 3.65 | 1.85 | 15.17 | 26.27 | 1.69 | 8,248 |
| **2013** | 2.45 | 33.15 | 5.69 | 0.02 | 3.95 | 13.14 | 0.58 | 2.12 | 2.84 | 2.86 | 14.55 | 17.35 | 1.29 | 9,335 |
| **2014** | 2.03 | 38.72 | 12.34 | 0.04 | 6.29 | 11.92 | 4.49 | 0 | 5.33 | 2.03 | 12.44 | 3.9 | 0.46 | 17,980 |
| **2015** | 2.43 | 23.2 | 0.92 | 0 | 4.51 | 5.09 | 7.11 | 0 | 5.82 | 3.12 | 32.91 | 13.34 | 1.56 | 13,686 |

| | | | | | | | | | | | | | | |
|---|---|---|---|---|---|---|---|---|---|---|---|---|---|---|
| **2016** | 12.23 | 28.76 | 5.73 | 0 | 7.85 | 7.68 | 3.14 | 1.01 | 2.9 | 6.61 | 11.49 | 5.31 | 7.3 | 10,825 |
| **2017** | 3.17 | 13.4 | 12.49 | 0 | 7.57 | 14.81 | 2.43 | 3.64 | 5.17 | 0.12 | 22.81 | 11.69 | 2.71 | 10,235 |
| **2018** | 1.56 | 24.42 | 13.06 | 0 | 11.22 | 9.44 | 8.2 | 1.26 | 2.58 | 0.15 | 14.73 | 13.32 | 0.07 | 9,626 |
| **2019** | 0 | 4.18 | 18.84 | 0 | 21.99 | 12.58 | 29.12 | 3.49 | 3.02 | 0.33 | 1.82 | 4.63 | 0 | 8,189 |
| **2020** | 0 | 0.05 | 16.44 | 0 | 75.29 | 0 | 0 | 0 | 0 | 2.63 | 0 | 0 | 5.59 | 2,165 |
| **2021** | 0 | 0 | 41.7 | 0 | 36.01 | 7.81 | 13.4 | 0 | 0 | 0 | 0 | 0 | 1.08 | 2,216 |
| **2022** | 0 | 13.71 | 17.55 | 0 | 68.75 | 0 | 0 | 0 | 0 | 0 | 0 | 0 | 0 | 1,459 |
| **2023** | 0 | 0 | 0 | 0 | 0 | 100 | 0 | 0 | 0 | 0 | 0 | 0 | 0 | 213 |

**Table S5.** Summary statistics from logistic regression analysis of ART-R transmission in prominent countries.

| Continent | Country | Year Range | Max Slope (%) - CI | Year of Max Slope | R2 | Years > 2% | First Year > 2% | Total Samples | Model Frequency for Last Observed Year | Raw Frequency for Last Observed year |
|---|---|---|---|---|---|---|---|---|---|---|
| Asia | Cambodia | 2002-2019 | 4.18 (2.33-6.03) | 2008 | 0.63 | 16 | 2002 | 6189 | 86.24 | 52.16 |
| Asia | Laos | 2010-2018 | 5.59 (2.95-8.23) | 2017 | 0.22 | 6 | 2012 | 2644 | 51.7 | 37.91 |
| Asia | Myanmar | 2004-2019 | 2.9 (1.58-4.22) | 2018 | 0.44 | 14 | 2004 | 11868 | 51.51 | 61.07 |
| Asia | Thailand | 1991-2017 | 6.33 (0-14.5) | 2012 | 0.78 | 16 | 1997 | 5599 | 75.38 | 67.5 |
| Asia | Vietnam | 2005-2019 | 5.45 (2.89-8.02) | 2013 | 0.64 | 13 | 2005 | 4557 | 75.78 | 75.81 |
| Africa | Eritrea | 2013-2019 | 5.5 (0-12.25) | 2019 | 0.93 | 2 | 2016 | 1548 | 15.67 | 15.3 |
| Africa | Rwanda | 2010-2023 | 3.25 (0-7.44) | 2023 | 0.89 | 4 | 2015 | 2153 | 18.03 | 17.37 |
| Africa | Sudan | 2014-2020 | 0.88 (0-2.06) | 2020 | 0.45 | 1 | 2020 | 1108 | 2.15 | 2.27 |
| Africa | Tanzania | 2003-2021 | 40.78 (0-118.81) | 2021 | 1 | 1 | 2021 | 3908 | 22.54 | 22.54 |
| Africa | Uganda | 1999-2022 | 9.12 (0-25.13) | 2022 | 0.96 | 6 | 2017 | 7752 | 28.82 | 25.72 |
| Oceania | Papua New Guinea | 1998-2020 | 3.08 (0-8.69) | 2020 | 0.53 | 2 | 2012 | 1401 | 6.32 | 6.38 |

**Table S6.** Summary of TES for artemisinin combination therapies in Africa that has reported delayed clearance rate or treatment failure rate above 10%.

| Reference | Year | Country, Site | Population | No. of samples with delayed clearance* | Delayed clearance (%) | No. of samples with treatment failure** | Treatment failure %(KM) | Association with K13 alleles and outcome | Association with *Pfmdr1* alleles and outcome |
|---|---|---|---|---|---|---|---|---|---|
| **Artemether-Lumefantrine** | | | | | | | | | |
| Plucinski et al., 2017 | 2015 | Angola, Zaire | Southern Africa | 1 | 1 | 8 | 11.9 | Not found | *pfmdr1* NYD and NFD haplotypes found in %91 of treatment failures |
| Dimbu et al., 2021 | 2019 | Angola, Lunda Sul | | 0 | 0 | 13 | 12.4 | Not found | N86 pfmdr1 allele found in 97% of treatment failures |
| Dimbu et al., 2024 | 2021 | Angola, Zaire | | 0 | 0 | 10 | 12 | Not tested | Not tested |
| Westercamp et al., 2022 | 2016 | Kenya, Siaya | Northeast Africa | 1 | 0.7 | 16 | 13.5 | Not found . | Not found |
| Gansane et al., 2021 | 2017 | Burkina Faso, Nanoro | West Africa | 1 | 0.9 | 22 | 26.5 | Not found | Not found |
| Moriarty et al., 2021 | 2018 | DRC, Mikalayi | Central Africa | 0 | 0 | 14 | 14 | Not found | Certain *pfmdr1* and *pfcrt* genotypes associated with treatment outcome. |

| | | | | | | | | | |
|---|---|---|---|---|---|---|---|---|---|
| Straimer et al., 2022 | | Rwanda, Kigali and Gasabo | | 5 | 27.8 | 1 | 5.6 | K13 alleles associated with delayed clearance. | Not tested |
| Ebond et al., 2021 | | Uganda, Busia | Northeast Africa | 0 | 0 | 10 | 12.8 | Not found | Not found |
| Ishengoma et al., 2024 | 2022 | Tanzania, Karagwe | East Africa | 11 | 12.5 | 1 | 1.7 | 561H mutation was significantly associated with delayed clearance. | Not found |
| **Artesunate-Amodiaquine** | | | | | | | | | |
| National Malaria Control Program*** | 2019 | Eritrea | Northeast Africa | 9 | 10.2 | 4 | 4.7 | Not tested | Not tested |
| Ishengoma et al., 2024 | 2022 | Tanzania, Karagwe | East Africa | 17 | 19.3 | 0 | 0 | 561H mutation was significantly associated with delayed clearance. | Not found |
| **Dihydroartemisinin–Piperaquine** | | | | | | | | | |
| Gansane et al., 2021 | 2018 | Burkina Faso, Gourcy | West Africa | 4 | 3.5 | 14 | 16.4 | Not found | Not found |
| | | Burkina Faso, Nanoro | | 2 | 1.8 | 9 | 11.3 | Not found | Not found |

*Delayed clearance has been defined as samples with parasitemia on day 3
**Treatment failure has been defined as PCR corrected efficacy based on Kaplan–Meier estimate.
***Data is obtained from Malaria Threats Map (WHO, 2024).

**Table S7. National adoption of first-line ACT regimens in different populations by the end of 2022.** The table shows the number of countries in each population that adopted each ACT regimen for the treatment of uncomplicated, unconfirmed, or confirmed *falciparum* malaria, according to the WHO (2023). Some countries have multiple first-line ACTs available for use and included in the table. The number of countries in each population is shown in parentheses. A: artemether, L: lumefantrine, AS: artesunate, AQ: amodiaquine, MQ: mefloquine, DHA: dihydroartemisinin, PYR: pyronaridine, CQ: chloroquine, PPQ: piperaquine, PQ: primaquine

| Population | AL | AS-AQ | DHA-PPQ | AS-PYR | AL+PQ | AS+SP+PQ | AS+MQ | AS+MQ +PQ | DHA-PPQ+PQ | AS-PYR +PQ |
|---|---|---|---|---|---|---|---|---|---|---|
| | Double ACTs | | | | Triple ACTs | | | | | |
| South America (9) | 2 | 0 | 0 | 0 | 5 | 0 | 0 | 1 | 0 | 0 |
| West Africa (18) | 16 | 1 | 0 | 0 | 1 | 0 | 0 | 1 | 0 | 0 |
| Central Africa (5) | 4 | 3 | 0 | 0 | 0 | 0 | 0 | 0 | 0 | 0 |
| East Africa (8) | 6 | 1 | 0 | 0 | 1 | 0 | 0 | 0 | 0 | 0 |
| Northeast Africa (7) | 3 | 2 | 0 | 0 | 2 | 0 | 0 | 0 | 0 | 0 |
| Southern Africa (5) | 5 | 1 | 0 | 0 | 0 | 0 | 0 | 0 | 0 | 0 |
| Western Asia (3) | 2 | 0 | 0 | 0 | 1 | 0 | 0 | 0 | 0 | 0 |
| Eastern South Asia (1) | 0 | 0 | 0 | 0 | 1 | 1 | 0 | 0 | 0 | 0 |
| Far-Eastern South Asia (2) | 1 | 0 | 0 | 0 | 2 | 0 | 0 | 0 | 0 | 0 |
| Eastern Southeast Asia (5) | 1 | 0 | 1 | 1 | 2 | 0 | 1 | 0 | 0 | 0 |
| Western Southeast Asia (2) | 0 | 0 | 0 | 0 | 1 | 0 | 0 | 0 | 1 | 1 |
| Oceania (4) | 2 | 0 | 1 | 0 | 1 | 0 | 0 | 0 | 0 | 0 |

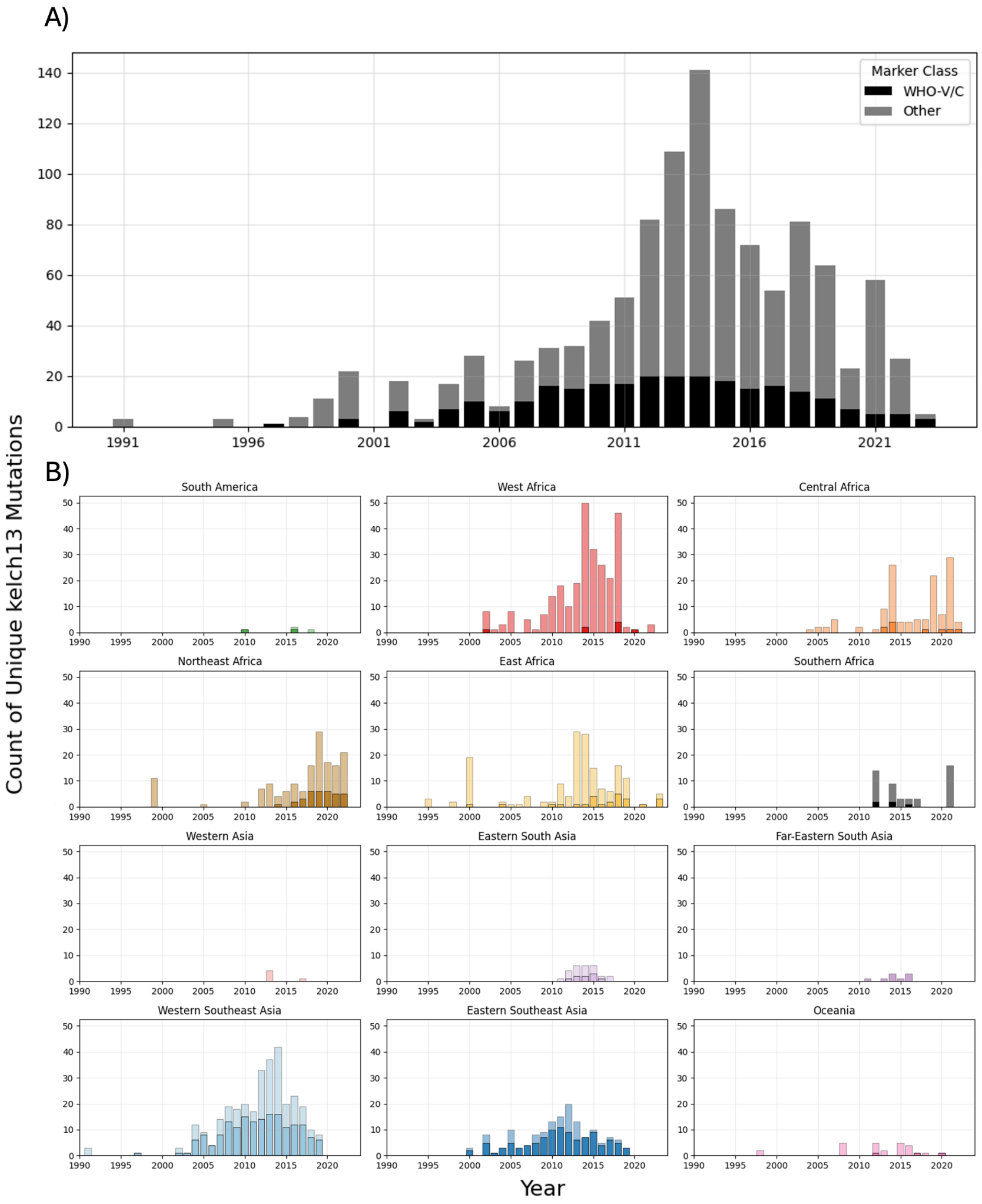


**Figure S1**. Number of unique WHO validated/candidate *kelch13* mutations and other propeller mutations over time. A) Global overview of unique mutations B) Unique mutations over time split by population. Darker hue = WHO-validated/candidate mutations, lighter hue = other propeller domain mutations. Bars represent stacked mutation counts. North Africa is not shown as no mutations were reported there.

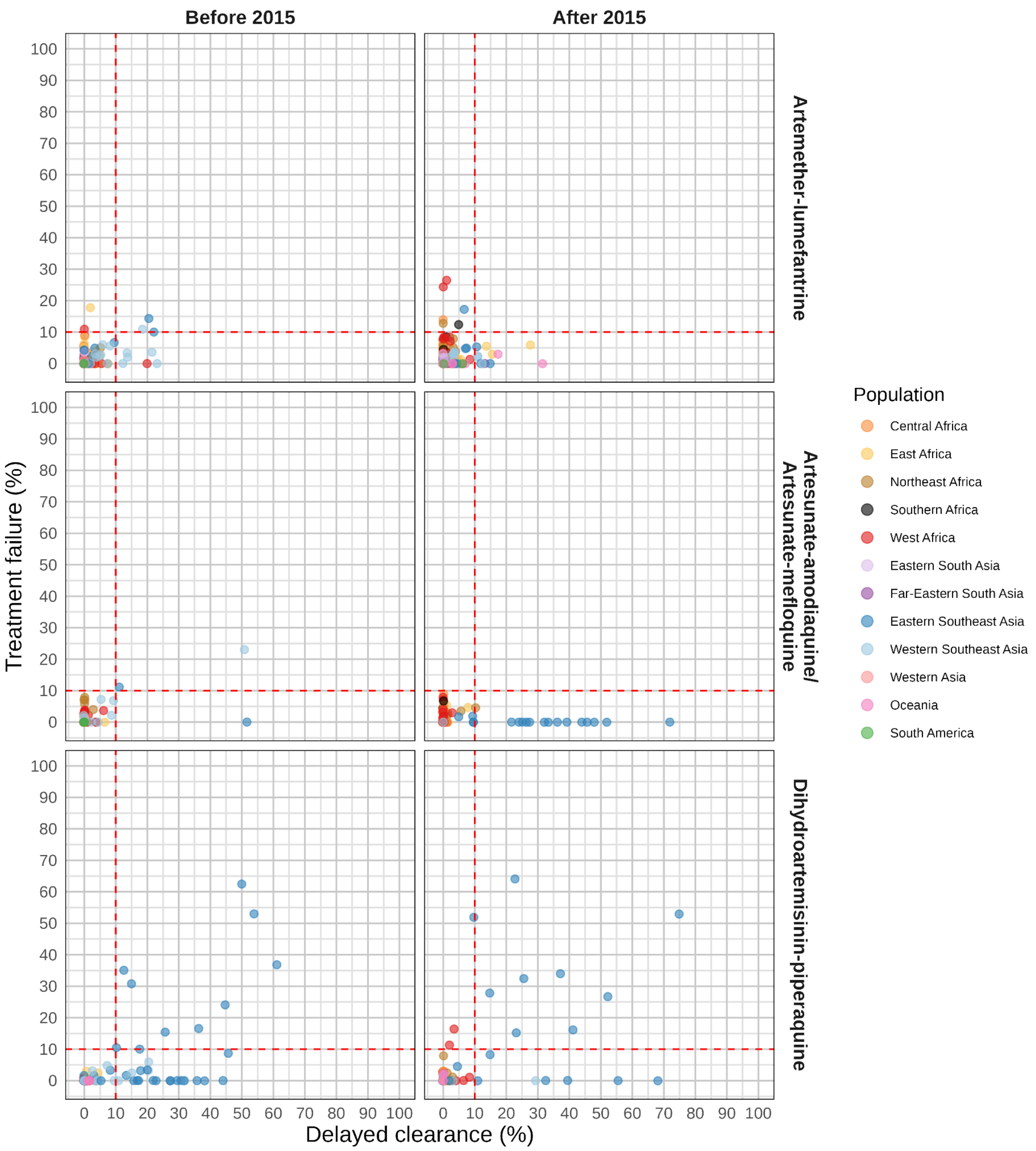


**Figure S2**. Scatter plot showing the results of TES monitoring 3 commonly used ACTs across populations, using the data from the WHO Malaria Threats Map (https://apps.who.int/malaria/maps/threats/). The x-axis represents delayed clearance, while the y-axis represents treatment failure rates. Scattered data points are colour-coded by region. Red dashed lines indicate the 10% threshold established by the World Health Organization (WHO) for policy change.

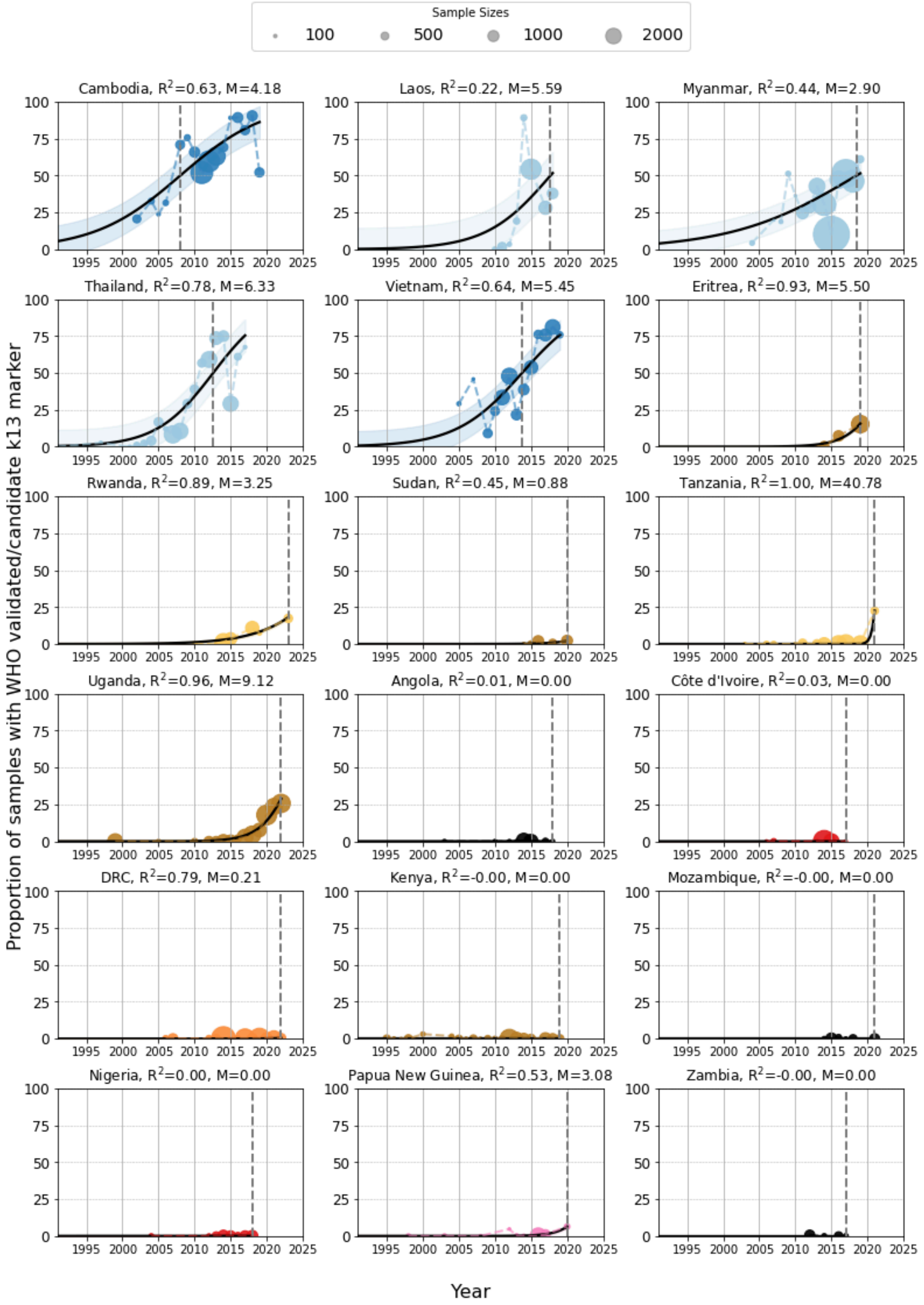


**Figure S3.** Percentage of samples with WHO validated/candidate *kelch13* propeller mutations over time. The observed data for each country is shown as a dashed line, coloured by population. The black line is the curve of a logistic model fitted to the observed data, which is forced to begin at 0 and end at 100%. The size of each point represents the number of samples collected in that country for each year. Timepoints with no line or value indicate years with no available data. Countries with fewer than three years with at least 25 samples each were excluded. Countries which showed no clear increase during this time were also excluded (Mali, Guyana, India, Ethiopia, China and Equatorial Guinea). The grey dashed line highlights the year in which the fitted curve reached its steepest point - i.e. the year with the fastest increase in frequency. The goodness-of-fit of the logistic curves varied substantially across countries ($R^2$ = 0.22-1.00). Notably, Tanzania had a very high growth rate, though this was based on only a single year where the frequency was above 2%.
