## Supplementary data README for "Understanding the Global Spread of Artemisinin Resistance: Insights from over 100K *Plasmodium falciparum* Samples"

**File: Supp_Dataset_S1.csv**

This file contains kelch13 marker per sample dataset from Balmer et al. (2024), released with the article.

**Data**

This file contains kelch13 marker and metadata for 112,933 samples. More details on data can be found in Methods section of the article.
Last Update: 07.08.2024

**Column Descriptions**

Each row represents a marker; each column represents a population described below.

| **Column Name** | **Description** |
|---|---|
| SampleId | Unique sample identification number. |
| Country | Country in which the sample was collected. |
| Population | Population of the country where sample was collected. These are SA (South America), AF-W (West Africa), AF-C (Central Africa), AF-NE (Northeastern Africa), AF-E (Eastern Africa), AS-S-E (Eastern South Asia), AS-F-FE (far-Eastern South Asia), AS-SE-W (Western Southeast Asia), AS-SE-E (Eastern Southeast Asia), and OC (Oceania). |
| Year | Year of collection for this sample. |
| Marker | Nonsynonymous amino acid change detected in kelch13 BTB/POZ and propeller domains at codon positions 349-726. |
| Continent | Continent where the sample was collected. |
| Source | Source of the data for this sample. This is either a publication, or Pf7 (MalariaGEN). |

**File: Supp_Dataset_S2.csv**

This file contains aggregated population data from Balmer et al. (2024), released with the article.

**Data**
This file contains count of kelch13 markers in 13 populations. Details on data can be found in Methods section of the article.
**Last Update:** 07.08.2024

**Column Descriptions**
Each row represents a marker; each column represents a population described below.

| Column Name | Description | |
|---|---|---|
| Marker | kelch13 nonsynonymous mutations detected. "3D7_REF" means no mutations. | |
| SA | Number of samples with the marker in this population: | South America |
| AF-W | | West Africa |
| AF-C | | Central Africa |
| AF-N | | North Africa |
| AF-NE | | Northeast Africa |
| AF-E | | East Africa |
| AF-S | | Southern Africa |
| AS-W | | Western Asia |
| AS-S-E | | Eastern South Asia |
| AS-S-FE | | Far-Eastern South Asia |
| AS-SE-W | | Western Southeast Asia |
| AS-SE-E | | Eastern Southeast Asia |
| OC | | Ocenia |
| Total | Total number of samples with the marker. | |
